# Acyl-CoA thioesterase-2 facilitates β-oxidation in glycolytic skeletal muscle in a lipid supply dependent manner

**DOI:** 10.1101/2023.06.27.546724

**Authors:** Carmen Bekeova, Ji In Han, Heli Xu, Evan Kerr, Brittney Blackburne, Shannon C. Lynch, Clementina Mesaros, Marta Murgia, Rajanikanth Vadigepalli, Joris Beld, Roberta Leonardi, Nathaniel W. Snyder, Erin L. Seifert

**Affiliations:** MitoCare Center; Department of Pathology and Genomic Medicine, Thomas Jefferson University, Philadelphia, PA 19107; Department of Biochemistry and Molecular Medicine, West Virginia University, Morgantown, West Virginia, 26506; Center of Excellence in Environmental Toxicology, Perelman School of Medicine, University of Pennsylvania, Philadelphia PA 19104; Max Planck Institute for Biochemistry, 82152 Martinsried, Germany; Department of Biomedical Sciences, University of Padova, 35131 Padua, Italy; Department of Microbiology and Immunology, Drexel University, Philadelphia, PA 19102; Lewis Katz School of Medicine, Temple University, Philadelphia, PA 19140

## Abstract

Acyl-Coenzyme A (acyl-CoA) thioesters are compartmentalized intermediates that participate in in multiple metabolic reactions within the mitochondrial matrix. The limited availability of free CoA (CoASH) in the matrix raises the question of how the local acyl-CoA concentration is regulated to prevent trapping of CoASH from overload of any specific substrate. Acyl-CoA thioesterase-2 (ACOT2) hydrolyzes long-chain acyl-CoAs to their constituent fatty acids and CoASH, and is the only mitochondrial matrix ACOT refractory to inhibition by CoASH. Thus, we reasoned that ACOT2 may constitutively regulate matrix acyl-CoA levels. *Acot2* deletion in murine skeletal muscle (SM) resulted in acyl-CoA build-up when lipid supply and energy demands were modest. When energy demand and pyruvate availability were elevated, lack of ACOT2 activity promoted glucose oxidation. This preference for glucose over fatty acid oxidation was recapitulated in C2C12 myotubes with acute depletion of *Acot2*, and overt inhibition of β-oxidation was demonstrated in isolated mitochondria from *Acot2*-depleted glycolytic SM. In mice fed a high fat diet, ACOT2 enabled the accretion of acyl-CoAs and ceramide derivatives in glycolytic SM, and this was associated with worse glucose homeostasis compared to when ACOT2 was absent. These observations suggest that ACOT2 supports CoASH availability to facilitate β-oxidation in glycolytic SM when lipid supply is modest. However, when lipid supply is high, ACOT2 enables acyl-CoA and lipid accumulation, CoASH sequestration, and poor glucose homeostasis. Thus, ACOT2 regulates matrix acyl-CoA concentration in glycolytic muscle, and its impact depends on lipid supply.

## INTRODUCTION

The mitochondrial matrix is replete with acyl-Coenzyme A (acyl-CoA) thioesters that are substrates and products of several reactions, including β-oxidation, branched chain amino acid catabolism, mitochondrial fatty acid synthesis (mtFAS), phospholipid hydrolysis, and covalent acylation reactions. The extent to which matrix acyl-CoA concentrations are regulated is poorly understood, but is metabolically important since CoA availability might become limiting (Abegaz, Martines et al., 2021, Himms-Hagen & Harper, 2001), long-chain acyl-CoAs inhibit the activity of some matrix proteins (Pande & Blanchaer, 1971, Paulson & Shug, 1984, Seiler, Martin et al., 2014, Woldegiorgis & Shrago, 1979, Woldegiorgis, Yousufzai et al., 1982), and mitochondrial lipid overload is associated with dysregulated glucose metabolism (Anderson, Lustig et al., 2009, Finck, Bernal-Mizrachi et al., 2005, Gavin, Ernst et al., 2018, Koves, Ussher et al., 2008, Muoio & Neufer, 2012, Smith, Lin et al., 2021). Mitochondrial acyl-CoA thioesterases (ACOT) hydrolyze acyl-CoA species of different chain lengths to their constituent fatty acids and CoASH (Bekeova, Anderson-Pullinger et al., 2019, Tillander, Alexson et al., 2017) and thus are candidates to regulate matrix acyl-CoA levels.

ACOTs reside in the cytosol, peroxisomes, and in mitochondria. Five ACOTs associate with mitochondria: ACOT2, ACOT7, ACOT9, ACOT11, and ACOT13; all localize to the matrix (Bekeova et al., 2019), although Acot13 appears to also associate with the outer mitochondrial membrane (Tillander et al., 2017). Tissue distribution of ACOTs varies by family member. ACOT7 is expressed mainly in the central nervous system (CNS) and ACOT11 is expressed in brown adipose tissue. ACOT2, ACOT9 and ACOT13 are widely expressed outside of the CNS, often co-expressed in the same tissue, and, together, cover a wide range of acyl-CoA chain length (Bekeova et al., 2019, Tillander, Arvidsson Nordstrom et al., 2014). What distinguishes these three Acots is the sensitivity of their activity to the matrix environment: ACOT9 and ACOT13 activity is strongly inhibited by CoA, whereas ACOT2 activity is insensitive to CoA and to other potential regulators that are present in the matrix (Bekeova et al., 2019, Tillander et al., 2014). Thus, ACOT2 may constitutively regulate acyl-CoA levels in the matrix.

The major substrates of ACOT2 are long-chain fatty acyl CoAs, with the highest affinity for C14:0- and C16:0-CoA. Affinity is similar to that of long- and medium-chain acyl-CoA dehydrogenase (Acadl, Acadm; the first step of β-oxidation) for those substrates. Thus, Acot2 is well-poised to regulate β-oxidation. Studies of ACOT2 overexpression in hepatocytes supported this potential role of ACOT2 in the regulation of β-oxidation, and also suggested that the long-chain fatty acids released by long-chain acyl-CoA hydrolysis could increase proton leak, presumably via fatty acid cycling within the inner mitochondrial membrane (Moffat, Bhatia et al., 2014). However, the role of endogenous ACOT2 has not been explored, and was the goal of this study. While the liver expresses low levels of ACOT2 (Bekeova et al., 2019), other organs have high expression, including skeletal muscle (SM) and heart. We reasoned that SM would be an interesting context to examine ACOT2-linked metabolic function because ATP demand and β-oxidation flux vary substantially (e.g., with fiber type, time of day, diet), allowing ACOT2 to be studied across a range of bioenergetic conditions.

Using mice with striated muscle-specific knockout of *Acot2* (Acot2-/-), we found that, *in vivo*, the respiratory quotient of Acot2-/- mice shifted towards even higher glucose oxidation at night. This shift was observed over a wide range of lipid supply. When mice were fed a normal chow diet or fasted overnight, the shift to higher glucose oxidation *in vivo* in Acot2-/- mice was accompanied by evidence for lipid overload in white quadriceps. This metabotype was evidenced by greater levels of acyl-CoAs including β-oxidation intermediates and by loss of correlation between metabolites of adjacent reactions of β-oxidation. This “β-oxidation overload” was mitigated when fasted Acot2-/- mice ran on a treadmill. When mice were fed a high fat diet, the shift to glucose oxidation in Acot2-/- mice was accompanied by lower levels of acyl-CoAs and β-oxidation intermediates, and better correlation between adjacent β-oxidation reactions in quadriceps. High-fat fed Acot2-/- mice showed improved whole-body glucose homeostasis, and lower levels of ceramides and their derivatives in white SM. Altogether, these observations support a model in which ACOT2 facilitates the oxidation of fatty acids in white SM when energy demand and lipid supply are low-to-modest. However, when lipid supply is high, ACOT2 promotes the accumulation of ceramide derivatives in white SM, conditions that could eventually counter glucose disposal. Thus, suppressing ACOT2 activity might be a strategy to defend glucose homeostasis when lipid supply is high.

## RESULTS

### ACOT2 expression relative to β-oxidation proteins is fiber type dependent

Because the abundance of enzymes and transporters related to substrate metabolism differs in different SM fiber types (Murgia, Nagaraj et al., 2015), and Acot2 (previously known as MTE1) has been linked to mitochondrial metabolism (Cole, Murray et al., 2011, Himms-Hagen & Harper, 2001, King, Young et al., 2007, Moffat et al., 2014), we first determined the fiber type-specific expression of Acot2 alone, and in relation to the major metabolic pathways that it is hypothesized to influence. To this end, we mined a mass spectrometry-based proteomics resource for mouse single SM fibers (Type 1, 2a, 2×, 2b) obtained from soleus and extensor digitorum longus muscles (Murgia et al., 2015). We first estimated Acot2 protein abundance, alone, and relative to oxidative phosphorylation (oxphos) capacity (sum of expression of ETC Complex I, III, IV and ATP Synthase subunits) and β-oxidation capacity (sum of levels of β-oxidation proteins). Acot2 expression was highest in Type 1 > Type 2a ≅ Type 2× ≅ Type 2b, whereas its expression relative to oxphos or β-oxidation capacity was Type 1 > 2a > 2b ≅ 2× (Fig1A). ACOT2 potentially competes with long-chain acyl-CoA dehydrogenase (ACADL) for substrate (see Fig4A); ACOT2 expression relative to that of ACADL was highest in Type 1 > 2a > 2b ≅ 2× (Fig1A). To obtain further insight into the potential relevance of ACOT2 in different fiber types, we evaluated 1) the capacity for β-oxidation relative to that for oxphos, 2) expression of ACADL relative to that of the upstream long-chain keto acyl-CoA thiolase (HADHB), and 3) expression of ACADL relative to that of upstream CPT2 that re-esterifies long-chain acyl-carnitines to the CoA form. For each of these three comparisons, expression in Type 2× and 2b exceeded expression in Type 1 or 2a (Fig1A). Thus, ACOT2 is least abundant in highly glycolytic Type 2b and 2× fibers, and ACOT2 and/or ACADL in Type 2b and 2× fibers *vs*. Type I or 2a fibers potentially receive the greatest inputs from CPT2 and/or HADHB.

**Fig. 1:**
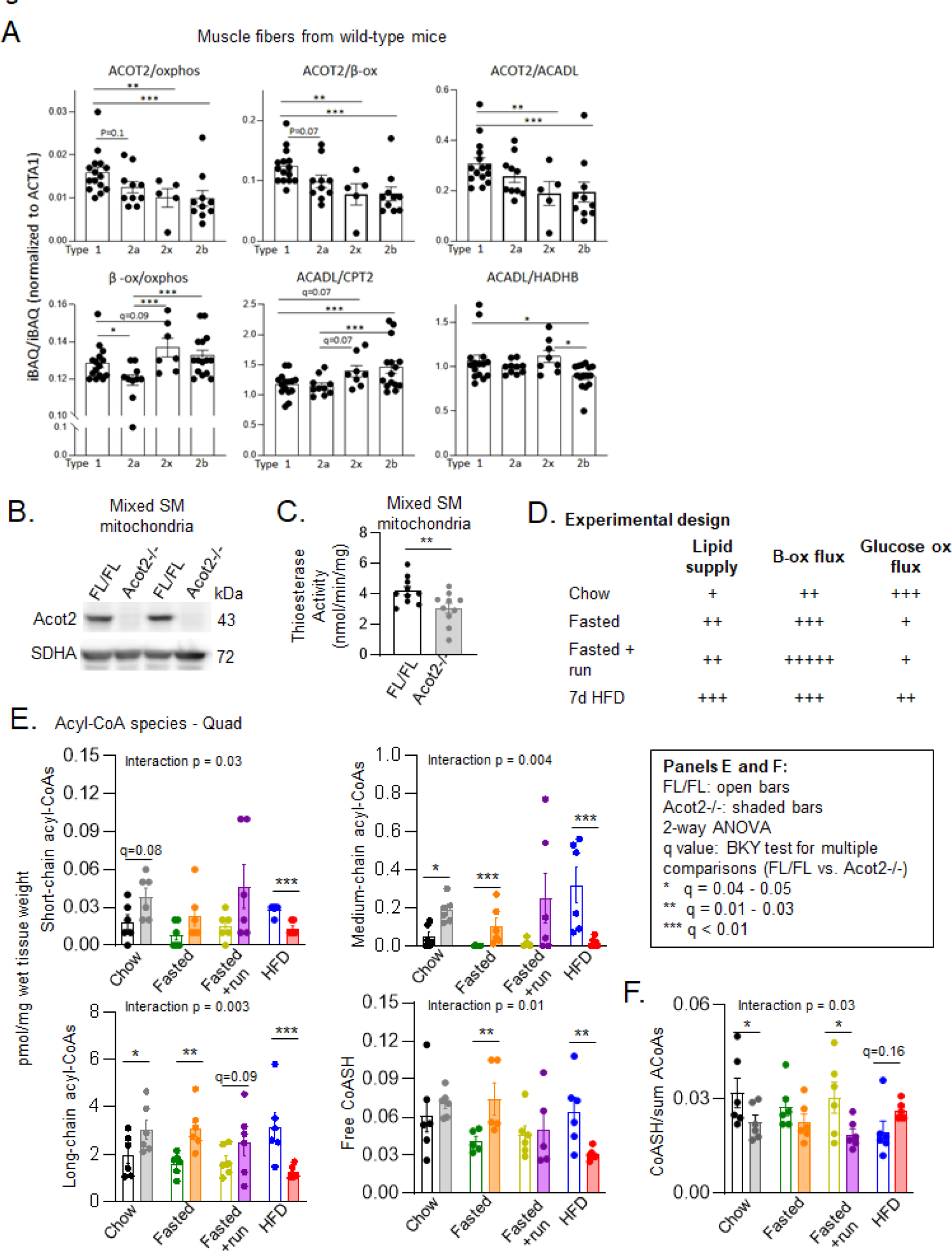
Model of Acot2 depletion in muscle and study design. A. Proteomics analysis from single murine muscle fibers. One ANOVA and multiple comparisons with correction for FDR (0.05; Benjamini, Krieger, Yekutieli method): * 0.05 ≤ q ≤ 0.03, ** 0.01 ≤ q < 0.03, *** q<0.01. B. Representative immunoblots comparing Acot2 levels in skeletal muscle (SM) mitochondria from FL/FL (Loxp/Loxp control) and Acot2-/- (knockdown of Acot2 in striated muscle) mice. SDHA (succinate dehydrogenase) was used as the loading control. C. Thioesterase activity measured in 40 µg of FL/FL and Acot2-/- SM mitochondria per reaction with 10 µM malonyl-CoA as substrate in n=10/genotype.** p=0.02, unpaired t-test. D. Summary of the experimental design. E. Short-, medium-, long-chain acyl CoAs and CoASH detected by high resolution mass spectrometry analysis of quadriceps muscle from FL/FL and Acot2-/- mice studied under chow, fasting, running after fasting, and HFD conditions, n=6/genotype. Statistics: two-way ANOVA and multiple compariFsons (q values) with correction for FDR (0.05; Benjamini, Krieger, Yekutieli method): * 0.05 ≤ q ≤ 0.04, ** 0.01 ≤ q < 0.03, *** q<0.01. N.S.: not significant. F. Ratio of CoASH to the sum of ACoAs (data from panel E). Statistics as in Panel E. Panels A, C, E, F: Bars are mean ± s.e.m, individual points are data from each mouse.

### ACOT2 plays a role in the β-oxidation pathway in SM

To investigate the role of endogenous ACOT2 in SM, we used a mouse model with Acot2 depletion in striated muscle (Acot2-/-), driven by the muscle creatine kinase promoter (CKM). The largest SM groups in the mouse comprise mainly Type 2× and 2b fibers. Thus, we focused on the white portion of Quadriceps (Quad) muscle, isolated mitochondria from the white portion of Quad, and also isolated mitochondria from all limb muscles (“mixed SM”). Acot2 was undetectable by western blot analysis of mixed SM mitochondria from Acot2-/- mice (Fig1B). Thioesterase activity for myristoyl-CoA (C14:0-CoA) was significantly lower in mixed SM mitochondria; residual thioesterase activity likely reflects that from Acot13 (Fig1C).

We reasoned that, given the diversity of possible processes that Acot2 could impact, a broad survey approach would allow us to unveil Acot2’s role in SM. The experimental design is summarized in Fig1D; we used diet, fasting, treadmill running of fasted mice, and light vs. dark phase of the day (rest *vs*. active phase, in mice) to generate different levels of lipid supply and energy demand. The intention of a short, 7-day, 60% HFD was to increase lipid supply substantially while avoiding many deleterious effects of a longer exposure. We note that transcript and protein levels of ACOT9 and ACOT13, two ACOTs expressed in SM and sensitive to CoA inhibition (Bekeova et al., 2019, Tillander et al., 2014)) were similar between Loxp and Acot2-/- Quad in all conditions, as were transcript levels of cytosolic and peroxisomal Acots (Suppl Fig1AB).

To obtain an overview of metabolism in the mitochondria matrix, we measured (in Quad) steady state levels of metabolites produced in the mitochondrial matrix, namely acyl-CoAs (ACoAs: β-oxidation), organic acids (Krebs cycle) and a subset of amino acids including branched chain amino acids (BCAA) whose metabolism, as for fatty acids, is heavily CoA-dependent. Metabolites were measured for all 4 conditions. Only ACoAs showed changes in all 4 conditions (Fig1E). In contrast, organic acids were not affected by Acot2 loss in any condition (Suppl Fig1C), and amino acids (Leu, Val, Gly, Gln) were slightly higher (∼30%) in Chow Acot2-/-, and Val was modestly (∼20%) lower in Fast+run Acot2-/-, but unaffected in Fast and HFD by *Acot2* loss (Suppl Fig1D). These findings support a primary role for ACOT2 in β-oxidation. The results also suggest that ACOT2 is not required for lipoic acid synthesis by the mtFAS pathway since otherwise levels of Krebs cycle metabolites and BCAAs would likely be altered.

Using liquid chromatography-high resolution mass spectrometry (LC-HRMS), we obtained absolute levels of the major ACoA species and CoASH. Samples were analyzed as a single batch, thereby facilitating comparisons between genotypes and across conditions (Fig1E). *Acot2* loss was associated with higher ACoA levels for all chain lengths in Chow, without a change in CoASH (Fig1E, black and grey bars). A similar pattern was evident in Fasted, though only medium- and long-chain ACoAs were higher in Acot2-/- Quad, and CoASH was also higher (Fig1E, green and orange bars). When Fasted mice ran on a treadmill (Fast+run), differences between Loxp and Acot2-/- largely disappeared and we note that CoASH was similar between genotypes (Fig1E, gold and purple bars). Short term HFD was associated with a rise in ACoAs of all chain lengths in Loxp Quad, but this was not apparent in Acot2-/- Quad, such that levels of ACoAs were lower in Quad of HFD Acot2-/- mice *vs*. Loxp mice (Fig1E, blue and red bars). CoASH was also lower in Quad from HFD-fed Acot2-/- (Fig1E, blue and red bars). An independent analysis further supported these changes with HFD. Using different cohorts of Loxp and Acot2-/- mice, total CoA (free and bound) was measured by HPLC in hydrolyzed Quad samples and was significantly lower in Quad from HFD-fed Acot2-/- mice (Suppl Fig2).

To estimate if CoASH availability changed in Acot2-/- Quad, we calculated the ratio of CoASH to the sum of ACoAs (short-, medium- and long-chain). Chow and also Fast+run were associated with significantly decreased ratios in Acot2-/- Quad *vs*. Loxp Quad, whereas Fasted was unchanged and HFD tended to be higher (Fig1F). Thus CoA sequestration in Quad (i.e., increased AcoAs) from Chow-fed, Fasted and Fast+run Acot2-/- mice was associated with lower CoASH availability in the Chow and Fast+run conditions. In Quad from HFD-fed Acot2-/- mice, CoASH availability tended to be higher despite the lower CoASH level.

### The impact of ACOT2 on β-oxidation does not depend on changes in protein abundance

We considered whether changes in the capacity of metabolic pathways might underlie changes in ACoA levels. To this end, RNAseq analysis was performed in Quad from mice in Chow, Fasted and HFD conditions. A subset of transcripts is presented in Fig2A. Volcano plots and pathway analyses are presented in Suppl Fig3. Full data sets are available (GEO accession #: GSE234495). In both Chow and Fasted conditions, Acot2 loss had little impact on transcript levels, including those associated with fatty acid uptake into SM and mitochondria; β-oxidation and peroxisomal fatty acid metabolism; CoA biosynthesis, mitochondrial transport and degradation; and fatty acid binding proteins (Fig2ABC). Transcript levels of ETC subunits were also unaffected. All the latter was also the case in HFD (Fig2ABC), though Fatty Acid Binding Protein 4 was elevated in Acot2-/- Quad (Fig2C), and small changes were evident in pathways not directly related to metabolism (Suppl Fig3). We note that there were no changes that would be consistent with the activation of the integrated stress response or mTORC1, previously associated with loss of CPT1b or Acyl-CoA synthetase 1 (Vandanmagsar, Warfel et al., 2016, Zhao, Pascual et al., 2019). Western blot analysis was performed for a subset of β-oxidation and CoA metabolism proteins in Quad from Chow- and HFD-fed mice, and showed no change with *Acot2* loss (Fig2D). These negative data suggest that changes in ACoAs in Acot2-/- Quad do not reflect changes in the capacity for fatty acid transport, β-oxidation, or CoA transport and metabolism, but likely reflect changes in flux.

**Fig. 2:**
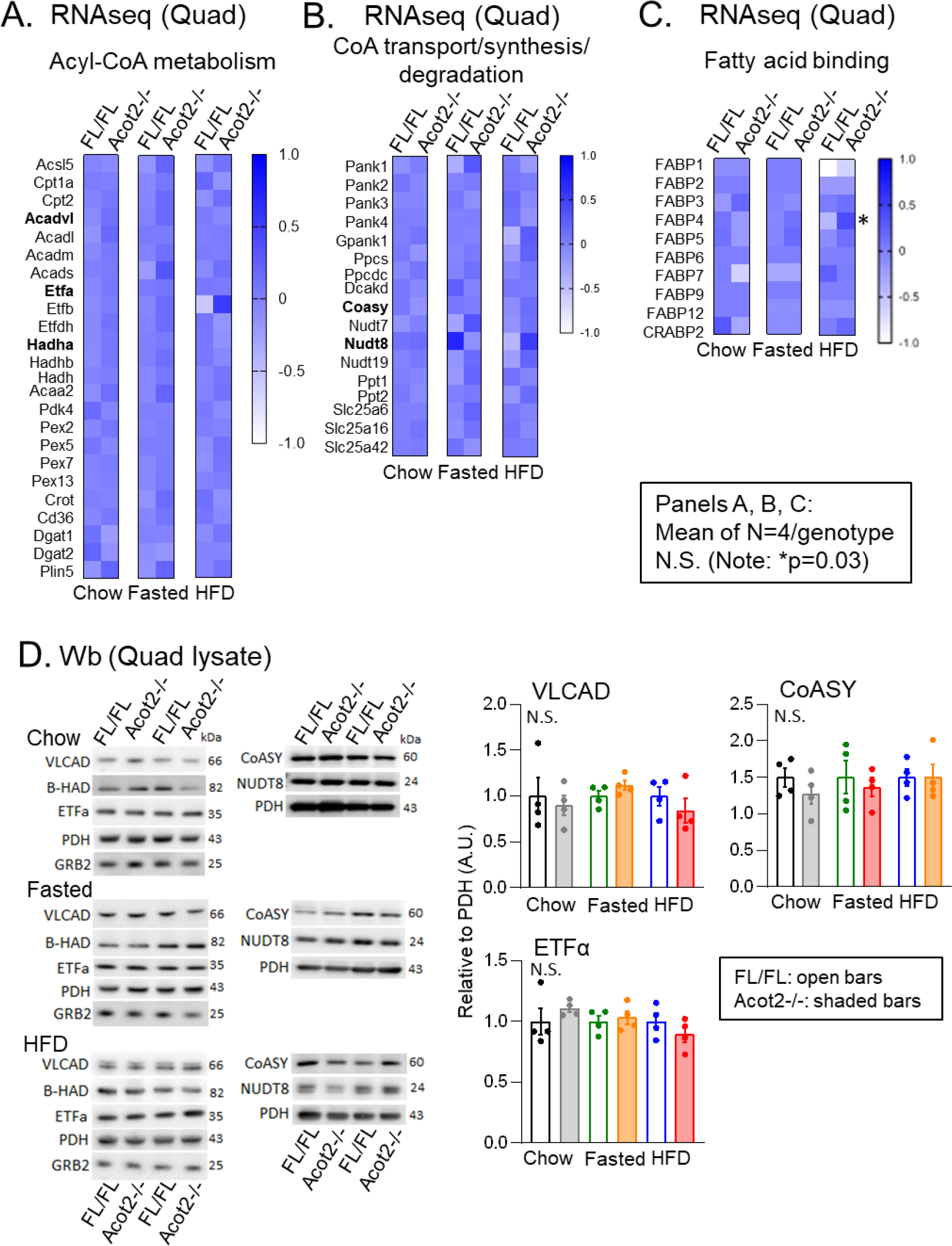
Lack of change, in any condition, in RNA or protein abundance of components of major CoA-related metabolic pathways in Quad from Acot2-/- mice. A-C. RNAseq analysis of Quad muscle, from mice fed normal chow, overnight fasted, or fed HFD for 7 days. Shown are transcript levels for enzymes in acyl-CoA metabolism (A), CoA synthesis/ degradation pathways (B), and fatty acid binding proteins (C), in Quad from chow-fed, overnight fasted and high fat diet (HFD)-fed mice. Each box is the average of n=4/genotype. D. Representative immunoblots from Quad lysates from chow-fed, fasted and HFD-fed mice, showing expression of VLCAD, β-HAD, ETFa, with PDH, GRB2 as loading controls; as well as CoASY and Nudt8, with PHD as loading control. At right: bar charts showing quantification. Values: mean ± s.e.m., points show individual mice; n=4/genotype/condition. N.S. Not significant (unpaired t-test).

### ACOT2 loss facilitates or limits β-oxidation in Quad mitochondria depending on lipid supply

To directly investigate how ACOT2 can alter β-oxidation in SM, bioenergetics analyses were performed using isolated SM mitochondria. Because the analysis of ACoA levels revealed condition-dependent genotype effects (Fig1E), analyses in mitochondria were conducted using different amounts of lipid substrate (palmitoylcarnitine (PCarn)), and over a range of simulated energy demand (achieved by varying in the ADP/ATP ratio using the creatine kinase (CK) clamp approach (Fisher-Wellman, Davidson et al., 2018)). PCarn oxidation generates palmitoyl-CoA and myristoyl-CoA that are substrates for ACOT2. As a first step, experiments were performed in mitochondria from mixed SM. Applying the CK clamp approach to mixed SM mitochondria from Chow- and HFD-fed mice, we could not detect any differences in O_2_ consumption rate (JO_2_) with Acot2 loss across a range of ADP/ATP ratio, whether at 20 µM or 40 µM PCarn (Suppl Fig4). We did not detect genotype differences in phosphorylating JO_2_ using saturating [ADP], and nor in maximal non-phosphorylating JO_2_ (using oligomycin to inhibit the ATP synthase) with 20 µM or 40 µM PCarn. Finally, there was no genotype difference for maximal ADP-stimulated or non-phosphorylating JO_2_ when pyruvate/malate was supplied to mitochondria (Suppl Fig4).

We then isolated mitochondria from white Quad and performed bioenergetics analyses. The CK clamp yielded a clear difference with Acot2 loss when we used 20 µM PCarn: JO_2_ tended to be higher at the highest ADP/ATP ratio (high energy demand), then fell more rapidly than Loxp mitochondria as ADP/ATP was decreased (Fig3A) such that the slope (“conductance”) was significantly steeper in Acot2-/- Quad mitochondria (Fig3B). Next, using saturating [ADP], we compared JO_2_ fueled by 20 *vs*. 40 µM PCarn (and 0.1 mM malate). We first noted that, with saturating [ADP], JO_2_ driven by 20 µM PCarn was significantly higher in Acot2-/- *vs*. Loxp mitochondria (Fig3C), consistent with the CK clamp experiment (Fig3AB). In contrast, JO_2_ with 40 µM PCarn was ∼36% lower than with 20 µM PCarn in Acot2-/- mitochondria, compared to only ∼19% lower in Loxp mitochondria (P = 0.04; Fig3C); this suggests that 40 µM PCarn led to lipid overload that was more pronounced in Acot2-/- mitochondria. In experiments using 20 µM PCarn and saturating [ADP], we noted that JO_2_ decreased over time; the drop was greater in Acot2-/- mitochondria (Fig3D), further supporting greater lipid overload. A similar analysis in mitochondria fueled with saturating pyruvate, malate and ADP showed little decrease over the same time interval (Fig3E). Also in the latter condition, JO_2_ was similar in Loxp and Acot2-/- mitochondria, and more than 2× the rate observed with 20 µM PCarn (Fig3E); thus genotype differences in PCarn-driven JO_2_ did not reflect a limitation of ETC capacity.

**Fig. 3:**
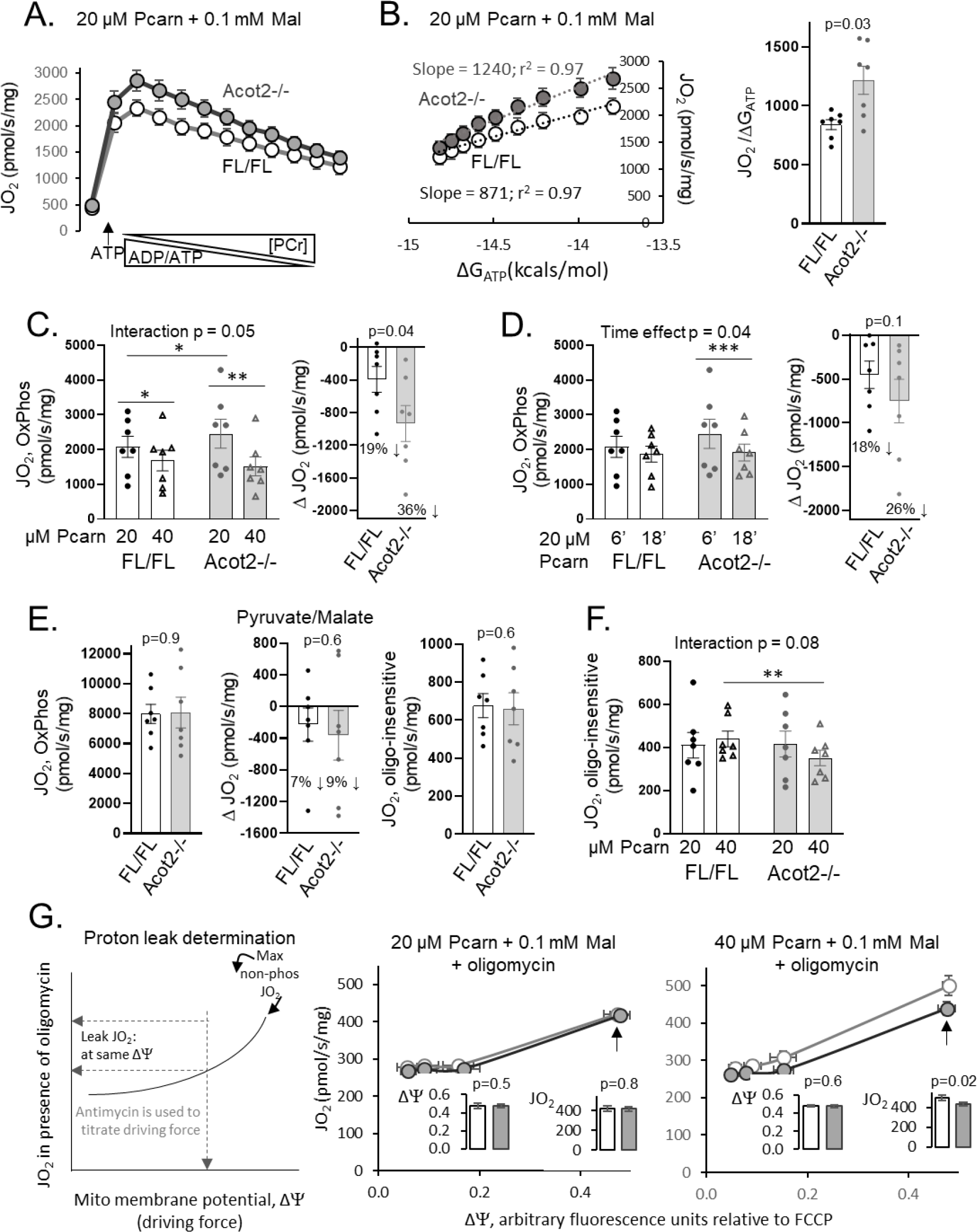
β-oxidation in mitochondria from white SM is stimulated or inhibited by Acot2 loss, depending on lipid supply. A. Oxygen consumption (JO_2_) in mitochondria from white SM, isolated from chow-fed, and supplied with 20 µM PCarn and 0.1 mM malate. The creatine kinase clamp approach was used to simulate changes in ATP demand (ADP/ATP) by varying phosphocreatine (PCr) concentration. Values: mean ± s.e.m.. B. The data from (A) were expressed as a function of Gibbs free energy for ATP (ΔG_ATP_). The slope reflects the conductance. Right panel: conductance; bar chart are mean ± s.e.m., individual points are from individual mice; p value: unpaired t-test, n=7/genotype. C. Left panel: JO_2_ measured in white SM mitochondria, with saturating [ADP] and 20 or 40 µM PCarn (plus 0.1 mM malate). Statistics: two-way ANOVA and multiple comparisons (q values) with correction for FDR (0.05; Benjamini, Krieger, Yekutieli method): * 0.05 ≤ q ≤ 0.03, ** 0.01 ≤ q < 0.03; n=7/genotype. Right panel: bar chart showing the difference in JO_2_ (ΔJO_2_) for 40 *vs*. 20 µM PCarn. p value: unpaired t-test, n=7/genotype. D. Left panel: JO_2_ measured in white SM mitochondria, with saturating [ADP] and 20 µM PCarn (plus 0.1 mM malate), showing measurements 6 and 18 minutes after ADP addition. Note: 6 minutes data for both genotypes are the same data shown in Panel C (20 µM PCarn) Statistics: two-way ANOVA and multiple comparisons (q values) with correction for FDR (0.05; Benjamini, Krieger, Yekutieli method): *** q < 0.01; n=7/genotype. Right panel: bar chart showing the difference in JO_2_ (ΔJO_2_) for 18 *vs*. 6 minutes; p value: unpaired t-test, n=7/genotype. E. Left panel: JO_2_ measured in white SM mitochondria, with saturating [ADP], 10 mM pyruvate and 5 mM malate (saturating [substrate]), measured 6 minutes after ADP addition. Middle panel: Difference in ADP-driven JO_2_ (ΔJO_2_) for 18 *vs*. 6 minutes Right panel: JO_2_ after addition of oligomycin to inhibit the ATP synthase. All panels: p value: unpaired t-test, n=7/genotype. F. JO_2_ measured in white SM mitochondria fueled by 20 or 40 µM PCarn (and 0.1 mM malate), after addition of oligomycin to inhibit the ATP synthase. Statistics: two-way ANOVA and multiple comparisons (q values) with correction for FDR (0.05; Benjamini, Krieger, Yekutieli method): ** 0.01 ≤ q < 0.03; n=7/genotype. G. Left panel: scheme illustrating the approach to evaluate proton leak in isolated mitochondria. Mitochondrial membrane potential (Δψm) and JO_2_ in the presence of oligomycin were measured in parallel in each preparation of mitochondria from white SM. Antimycin (inhibits Complex III of the ETC) was used to decrease the driving force. Middle panel: Proton leak measured in mitochondria fueled with 20 µM PCarn and 0.1 mM malate. Right panel: Same experiment as in Middle panel, except using 40 µM PCarn and 0.1 mM malate. Middle and right panels: insets show bar chart of mean (±s.e.m.) Δψm and JO_2_ at the arrow; p value: unpaired t-test, n=6/genotype. Panels B, C, D, E, F: Bar chart: mean ± s.e.m., individual points show data from each mice.

To gain insight into whether ACOT2 plays a role in non-phosphorylating JO_2_ (i.e., via fatty acid cycling causing proton leak) we measured maximal non-phosphorylating JO_2_ using oligomycin to inhibit the ATP synthase. With 20 µM PCarn, non-phosphorylating JO_2_ was similar between Loxp and Acot2-/- mitochondria. However, with 40 µM PCarn, non-phosphorylating JO_2_ was slightly lower in Acot2-/- *vs*. Loxp mitochondria (Fig3F). To directly evaluate proton leak, JO_2_ and membrane potential (delta psi; ΔΨm) were measured in parallel in the same mitochondrial preparation, in the presence of oligomycin and with titration of the electrochemical driving force using antimycin (Complex III inhibitor). By measuring non-phosphorylating JO_2_ at the same driving force (ΔΨm), a difference in non-phosphorylating JO_2_ indicates a difference in proton leak, (rather than a change in driving force) (see graphical explanation in Fig3G). With 20 µM PCarn, proton leak-driven JO_2_ was similar in both genotypes (Fig3G). With 40 µM PCarn, proton leak-driven JO_2_ was slightly but significantly lower in Acot2-/- mitochondria (Fig3G), indicating a greater proton leak in Loxp *vs*. Acot2-/- mitochondria fueled with 40 µM PCarn.

### Impact of ACOT2 on β-oxidation in white SM in vivo depending on lipid supply

To investigate how ACOT2 alters β-oxidation in SM *in vivo*, we used two approaches: analyses of steady state β-oxidation intermediates in Quad from mice in 4 conditions (Fig4) and whole-body energetics analyses by indirect calorimetry (IC) of Chow, Fasted and HFD mice (Fig5).

**Fig. 4:**
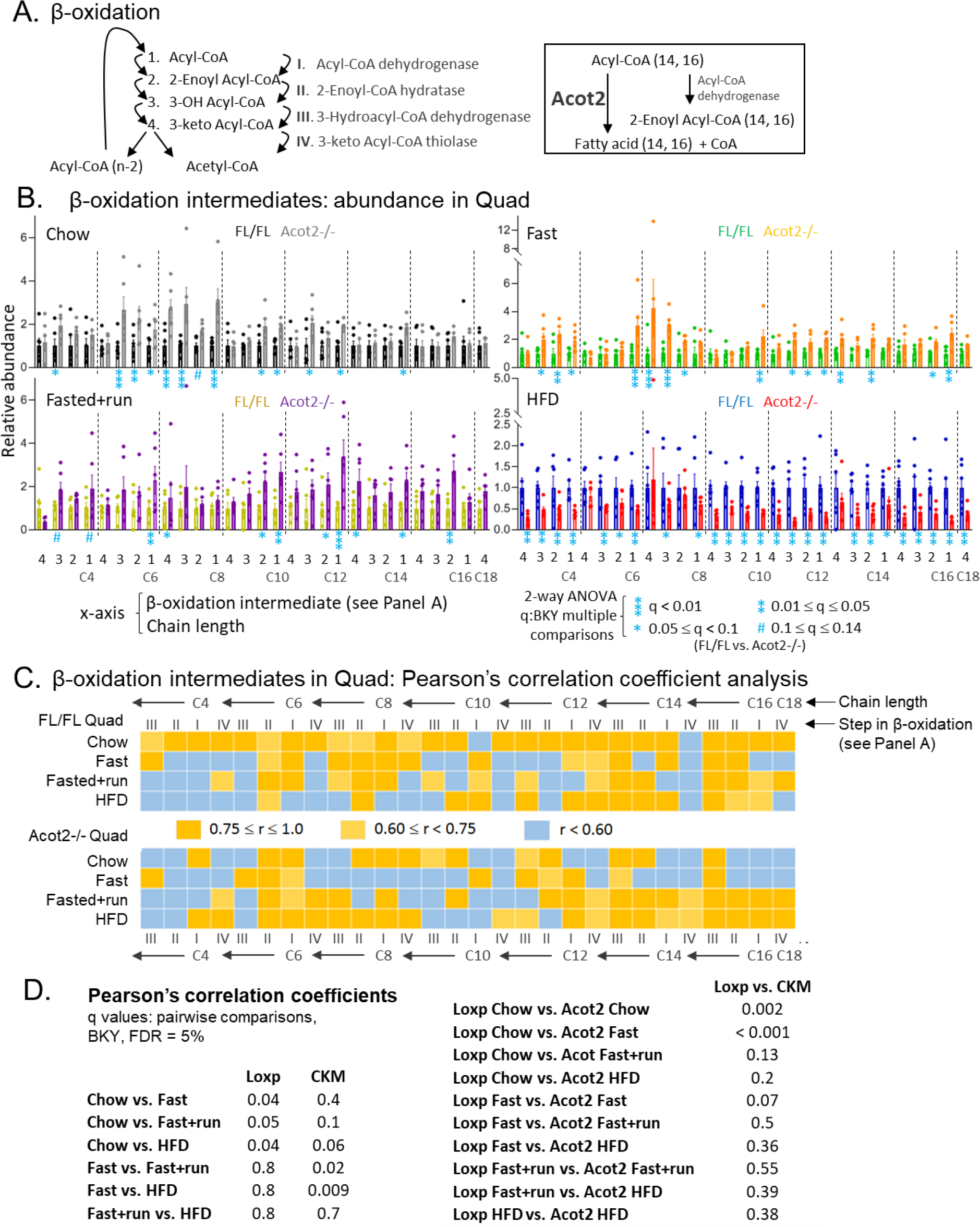
β-oxidation intermediates provide evidence for overload in white SM from Chow-fed and fasted Acot2-/- mice. A. Schematic representation of β-oxidation pathway and Acot2 reaction. B. Relative abundance of the β-oxidation intermediates for each reaction (1-4, corresponding to steps 1-4 shown in (A)) for all chain lengths (indicated by C4, C6 etc). Metabolites were measured by high resolution mass spectrometry. Statistics: shown within the panel; BKY: Benjamini, Krieger, Yekutieli, to correct for multiple comparisons, FDR: p=0.05. n=6/genotype/condition. C. Pearson’s correlation coefficient (r) analysis between adjacent steps of β-oxidation, using the intermediates shown in (B). Roman numerals: enzymes for each step (see panel A). D. Analysis of Pearson’s correlation coefficients. Statistics: shown within the panel; BKY: Benjamini, Krieger, Yekutieli, to correct for multiple comparisons, FDR: p=0.05.

**Fig. 5:**
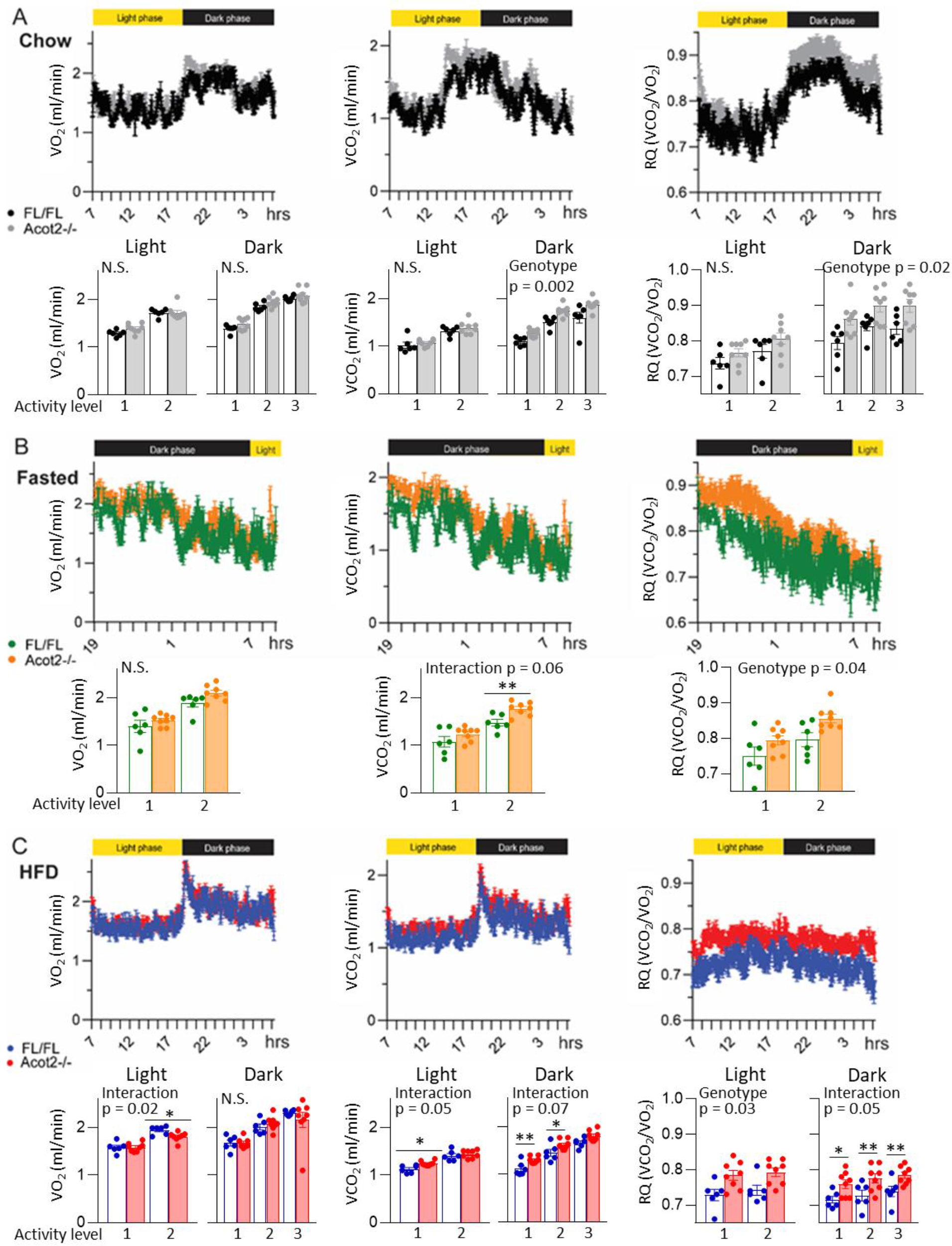
Indirect calorimetry shows a switch to greater glucose oxidation in Acot2-/- mice. A. Time course (time of the day) showing whole-body O_2_ consumption (VO_2_), CO_2_ production (VCO_2_) and respiratory quotient (RQ), during the light and dark phase of the day, in chow-fed mice. Bar chart shows VO_2_, VCO_2_ and RQ values in different activity bins (1 is lowest activity bin, 3 is the highest). For each mouse, data were averaged over 2 days, after a day of acclimation. B. Same as (A), but in overnight-fasted mice; food was removed at 7pm (19-h). C. Same as (A), but in mice fed 60% HFD for 7 days; data reflect the last 24 hrs of HFD. In all panels: Time course measurements show mean ± s.e.m. Bar charts show mean ± s.e.m., and individual points show data from each mice. Statistics: two-way ANOVA, Statistics: two-way ANOVA and multiple comparisons (q values) with correction for FDR (0.05; Benjamini, Krieger, Yekutieli method): * 0.05 ≤ q ≤ 0.03, ** 0.01 ≤ q < 0.03; n=6/genotype/condition.

Analyses of steady state levels of major ACoA species in Quad (Fig1E) also provided us with the levels of all β-oxidation intermediates (in relative value), for all 4 reactions of β-oxidation (Fig4A), for chain lengths from C18 down to C4, in all 4 conditions. In general, the amplitude of the intermediates (Fig4B) showed a similar pattern of change with *Acot2* loss as what we observed for the ACoAs (Fig1E, and in relative value in Fig4B). Specifically, for Chow and Fasted conditions in Acot2-/- Quad, while increases in the level of the intermediates were observed for most chain lengths, the most affected chain lengths were C6 and C8 (Fig4B). These increases for C6 and C8 intermediates were less apparent when fasted Acot2-/- mice ran on a treadmill (Fig4B). For the HFD condition, the abundance of intermediates was lower in Acot2-/- Quad, for all chain lengths (Fig4B), as was observed for ACoAs (Fig1E).

We further analyzed the β-oxidation intermediates by calculating Pearson’s correlation coefficients (r) between adjacent steps of β-oxidation (Fig4C). We reasoned that this, together with information about amplitudes, would provide insight into changes in flux *in vivo* (Ferrara, Wang et al., 2008, Steuer, 2006). In Fig4C, each square represents a correlation coefficient coded by color: dark orange for highly correlated (r ≥ 0.75), light orange for moderately correlated (0.6≤ r <0.75), blue for poorly correlated (r < 0.6) (Schober, Boer et al., 2018). Roman numerals indicate the step of β-oxidation (see Fig4A). As one approach to statistically compare conditions and genotypes, we performed 2-way ANOVA on the average r values for each comparison (i.e., adjacent reaction, all chain lengths), 4 conditions, and 2 genotype, and used a false discovery rate of 5% for post-hoc pairwise comparisons (Fig4D). We also calculated confidence intervals (65%) for the Pearson’s correlation coefficients (Suppl Fig5A), and performed statistics on the lower limit of the confidence intervals using a false discovery rate of 5% (Suppl Fig5B). High correlation is suggestive of first order kinetics, and low correlation indicates a loss of first order kinetics that can have several causes. In Quad from Chow-fed Loxp mice, β-oxidation was well-correlated for long-chain and short-chain species, and moderately correlated for medium-chain species, with only 2 poorly correlated nodes. Loxp Quads from the Fasted and Fasted+run conditions, and most strikingly the HFD condition, showed much less correlation between adjacent steps of β-oxidation as compared to Quad from Chow-fed Loxp mice, especially for medium- and short-chain species; there were 14, 12 and 15 poorly correlated nodes for Fasted, Fasted+run and HDF, respectively (Fig4C, “FL/FL”). With loss of ACOT2, β-oxidation in Quad from Chow-fed mice was poorly correlated, with 15 poorly correlated nodes mainly associated with long- and short-chain species. Poor correlation remained evident in Acot2-/- Quad from Fasted mice, with 18 poorly correlated nodes spread across all chain lengths. Treadmill running of fasted Acot2-/- mice led to an improvement in correlation (10 poorly correlated nodes for medium- and short-chain). Finally, in striking contrast to Quad from HFD-fed Loxp mice, Quad from HFD-fed Acot2-/- mice showed well-correlated β-oxidation, with only 7 poorly correlated nodes clustered mainly among the short-chain intermediates. Loss of correlation in Acot2-/- Quad from Chow and Fasted mice, combined with a greater abundance of some β-oxidation intermediates, supports lipid overload during β-oxidation in Acot2-/- Quad, *in vivo*, under those conditions. The analysis also suggests that treadmill running of Fasted Acot2-/- mice relieved lipid overload in Quad (but, nonetheless, sequestered CoASH to an extent that CoASH was lower (Fig1F), and that loss of Acot2 lessened the lipid overload in the HFD condition.

As a second approach to investigate Acot2’s role in β-oxidation in SM, *in vivo*, we used indirect calorimetry (IC) to evaluate *in vivo* substrate utilization in Chow, Fasted and HFD conditions; SM contributes substantially to the indirect calorimetry signal because it is a large (30-40%) fraction of body weight (Kummitha, Kalhan et al., 2014, Rolfe & Brown, 1997). There was no difference in body composition in any condition (Suppl Fig6); thus, IC data was expressed per mouse. Whole-body O_2_ consumption and CO_2_ production (VO_2_ and VCO_2_, respectively) showed the expected diurnal pattern in Chow-fed mice (e.g., (Moffat et al., 2014)) and blunted amplitude in HFD-fed mice (Coomans, van den Berg et al., 2013)(Fig5); thus mice were well-acclimated to the setup. To evaluate for possible energy demand differences, we binned the locomotor activity data into 3 levels (the highest activity bin was not always included during the light phase because of insufficient data points in some mice). Then, in each mouse, the corresponding VO_2_, VCO_2_ and respiratory quotient (RQ: VCO_2_/VO_2_) data for each activity bin were averaged. VO_2_ was similar between genotypes, except for a decrease in HFD-fed Acot2-/- mice associated with the highest activity bin in the light phase (Fig5, lowest panel). In contrast, Acot2-/- mice showed higher VCO_2_ in all conditions, though only during the dark phase of the day (in fasted mice, this was reflected in the higher activity bin) (Fig5). RQ was also higher in Acot2-/- mice at night, generally independent of activity level, indicating greater glucose oxidation (Fig5).

To gain mechanistic insight into the IC findings, we note the lack of major changes in transcript levels in Quad for proteins involved in β-oxidation or fatty acid handling (Fig2ABC), and nor in the protein abundance of several beta-oxidation enzymes that we queried (Fig2D). We also queried the RNAseq data to determine transcript abundance, in Quad, of glucose transporters and glycolysis enzymes. There were no changes that reached statistical significance using adjusted p values. However, with HFD, we noted slightly higher transcript levels for some glycolytic enzymes in Acot2-/- Quad (*Ldha*: p = 0.03, *Eno1*: p = 0.05, *Aldoa*: p = 0.15, *Gpd1*: p=0.16, PKM: p = 0.11; unpaired t-test), and lower levels for others (*Slc2a3*: p = 0.02; *Hk1*: p = 0.05; unpaired t-test) (Suppl Fig7A). Finally, we tested, in Quad, well-established regulatory mechanisms that bias towards β-oxidation or glucose oxidation, namely pyruvate dehydrogenase phosphorylation (inhibits β-oxidation), levels of malonyl-CoA (inhibits CPT1) and acetyl-CoA carboxylase phosphorylation (inhibits malonyl-CoA synthesis). ACOT2 loss did not alter any of the latter regulatory mechanisms under Chow, Fasting or HFD (Suppl Fig7BCD). Because malonyl CoA levels were very low, we tested levels in mice that were fasted (24 hrs) then refed (6 hrs); malonyl-CoA levels in Quad tripled, but were independent of Acot2 expression (Suppl Fig7D).

To determine if the higher glucose oxidation in Acot2-/- mice was cell-autonomous, *Acot2* was acutely depleted (Acot2 kd) from C2C12 myotubes using different two siRNA duplexes (Suppl Fig8A). Myotubes (6 days after serum withdrawal) were incubated with glucose (10 mM) and palmitate:BSA (6:1, 100 µM palmitate, plus 0.5 mM carnitine). We evaluated the JO_2_ sensitive to mitochondrial pyruvate transporter inhibition by UK5099 (2 µM) (Yang, Ko et al., 2014). Acot2 kd resulted in higher JO_2_ and greater UK5099-inhibitable JO_2_ (Suppl Fig8BD), without a change in total capacity for glucose oxidation, determined using the chemical uncoupler FCCP (Carbonyl cyanide 4-(trifluoromethoxy)phenylhydrazone) (Suppl Fig8BD). To evaluate glycolysis, we measured the extracellular acidification rate (ECAR) sensitive to 2-deoxyglucose (2-DG). 2-DG-sensitive ECAR and maximal 2-DG-sensitive ECAR (in presence of FCCP) tended to be higher with Acot2 kd (Suppl Fig8CE).

We also evaluated β-oxidation in C2C12 myotubes with acute Acot2 kd, as JO_2_ sensitive to the CPT1 inhibitor etomoxir (4 µM) (Divakaruni, Hsieh et al., 2018). To increase reliance on fatty acids for ATP production, myotubes were cultured in substrate-limited media overnight, prior to measurements (see Methods). Recordings were performed with glucose (10 mM) and palmitate:BSA (6:1, 100 µM palmitate, plus carnitine). Prior to etomoxir addition, Acot2 kd myotubes had higher JO_2_. Etomoxir lowered the JO_2_ to a similar extent in control and kd cells (Suppl Fig8FH). β-oxidation capacity was also similar, determined using FCCP (Suppl Fig8FH). Prior to etomoxir, 2-DG-sensitive ECAR was higher in Acot2 kd myotubes. Etomoxir increased the ECAR in control and kd cells, but more so in the control (Suppl Fig8GI). These data suggest that β-oxidation was hindered in Acot2 kd cells, necessitating greater glycolysis.

### Lower levels of ceramides and derivatives in Quad from HFD-fed Acot2-/- mice and improved glucose homeostasis in Acot2-/- mice

The shift away from β-oxidation *in vivo* revealed by the IC experiments (Fig5) raised the question of the fate of fatty acids in Acot2-/- muscle. To this end, we evaluated lipids in white Quad. We note first that WAT weight was not different in any of the conditions (Suppl Fig6). Targeted lipidomics analysis by LC-HRMS was performed on extracts from Quad harvested from chow- and HFD-fed mice. Figure 6A shows the total abundance for each species. Acyl chain-specific results are shown in Suppl Table 1. In Quad from chow-fed mice, there was no genotype difference for any of the lipid species (Fig6A; Suppl Table 1). HFD-feeding led to an increase in diacyl- and triacylglycerides, of similar magnitude, in Quad from Loxp and Acot2-/- mice (Fig6A), without acyl chain-specific genotype effects (Fig6A; Suppl Table 1). Triacylglycerides were also assayed as total glycerol in hydrolyzed Quad samples, with the same result (Suppl Fig9). In Loxp Quads, HFD also led to robust increases in level of hexosylceramides and sphingomyelins, with no change in ceramides (Fig6A). In contrast, in Quads HFD-fed Acot2-/- mice, there were no changes in the level of hexosylceramides and sphingomyelins, and there was a drop in ceramide levels (Fig6A). Analysis of acyl chain-specific differences did not show a predominance for a particular acyl-chain species in the genotype effects (Suppl Table 1). Triacylglycerides were also measured biochemically in liver and heart, and were similar between genotypes, in both Chow and HFD conditions (Fig6B). In contrast, triacylglycerides and non-esterified fatty acid levels in serum were higher in HFD-fed Acot2-/- mice compared to HFD Loxp mice (Fig6B).

**Fig. 6:**
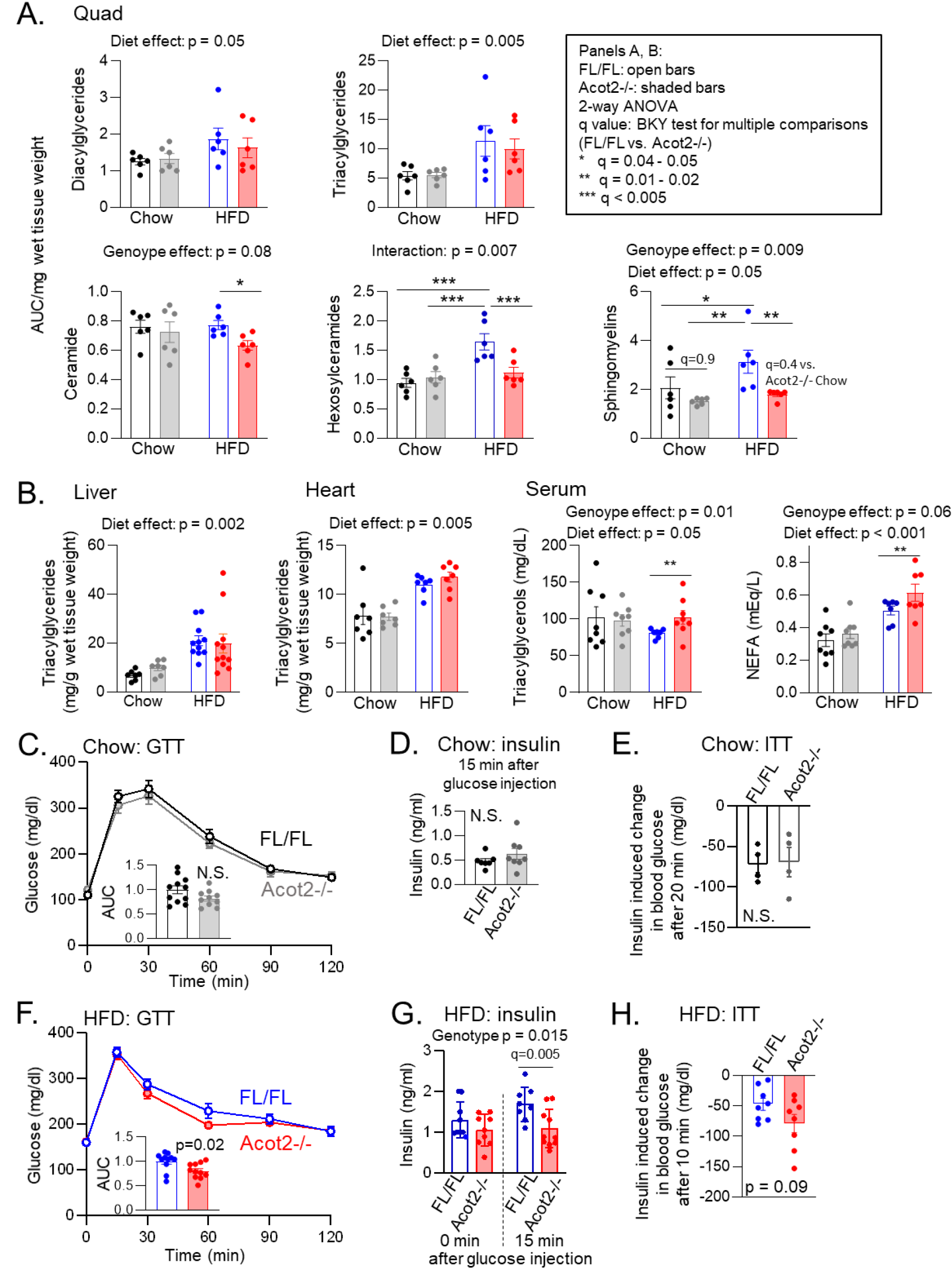
Less lipid accumulation in Quad and faster glucose disposal in HFD-fed Acot2-/- mice. A. Abundance of lipids, measured by high-resolution LC-MS analysis, in Quads from chow- and HFD-fed mice, n=6/genotype. Statistics are shown in the panel. BKY: Benjamini, Krieger, Yekutieli, to correct for multiple comparisons, FDR: p=0.05; * 0.05 ≤ q ≤ 0.03, ** 0.01 ≤ q < 0.03, *** q<0.01. B. Triacylglycerols measured in liver, heart and serum, measured biochemically, in chow and HFD fed mice, n=7-11/genotype. Statistics: two-way ANOVA and multiple comparisons (q values) with correction for FDR (0.05; Benjamini, Krieger, Yekutieli method): ** 0.01 ≤ q < 0.03. C. Oral glucose tolerance test (GTT, 2 µg/ 1 g of body weight glucose injection) in chow-fed mice, n=11 FL/FL and n=10 Acot2-/-. N.S.: not significant, unpaired t-test. D. Serum insulin levels 15 min after oral glucose (2 µg/ 1 g of body weight) injection in chow-fed mice, n=7 FL/FL and n=8 Acot2-/-. N.S.: not significant, unpaired t-test. E. Insulin tolerance test (ITT, 1.5 U/ kg of body weight insulin injection) in chow-fed mice, n=4/genotype. N.S.: not significant, unpaired t-test. F. Oral glucose tolerance test (GTT, 1 µg/ 1 g of body weight glucose injection) in HFD-fed mice, n=11/genotype. P value: unpaired t-test. G. Serum insulin levels before (in n=9/genotype) and 15min after (n=8 FL/FL, n=10 Acot2-/-) oral glucose (1 µg/ 1 g of body weight) injection in HFD-fed mice. Statistics: two-way ANOVA and multiple comparisons (q values) with correction for FDR (0.05; Benjamini, Krieger, Yekutieli method). H. Insulin tolerance test (ITT, 3 U/ kg of body weight insulin injection) in HFD-fed mice, n=9 FL/FL and n=8 Acot2-/-. P value: unpaired t-test. Panel A-F: values are mean ± s.e.m. The points represent individual experiments.

Finally, systemic glucose homeostasis can be influenced by mitochondrial β-oxidation (Anderson et al., 2009, Finck et al., 2005, Gavin et al., 2018, Koves et al., 2008, Muoio & Neufer, 2012, Smith et al., 2021), and insulin-mediated glucose uptake can be inhibited by ceramide and diacylglycerol (Petersen & Shulman, 2018). Thus we measured glucose clearance by oral glucose tolerance test (GTT: Fig6CF), insulin levels during the GTT (Fig6DG), and glucose levels after insulin injection (Fig6EH). For all parameters, Chow-fed Loxp and Acot2-/- mice were similar (Fig6CDE). In contrast, HFD-fed Acot2-/- mice showed faster glucose clearance (Fig6F) associated with lower insulin at 15 mins post glucose delivery (time 0 insulin was not different between genotypes)(Fig6G). Furthermore, after insulin injection, there was a tendency for a greater decrease in blood glucose in HFD-fed Acot2-/- mice (Fig6H).

## DISCUSSION

Four groups of observations support the role of endogenous ACOT2 in SM, especially in white SM. First, three independent approaches indicate that the primary role of ACOT2 is to promote β-oxidation. Indeed, in the absence of ACOT2, mitochondrial glucose oxidation was higher, *in vivo* and in C2C12 myotubes acutely depleted of ACOT2, and the switch to glucose oxidation was most evident when fatty acids and pyruvate were both readily available to be oxidized. Second, *in vivo*, Acot2 mitigated lipid overload in mitochondria from white SM when lipid supply and energy demand were low-to-modest. Alleviation of lipid overload by ACOT2 was also apparent in isolated mitochondria under high lipid supply. Third, in the context of HFD, ACOT2 favored the accumulation of ceramides and ceramide derivatives in white SM, along with slower glucose disposal. Finally, flux through ACOT2 led to higher proton leak in isolated mitochondria under high lipid supply; this may be relevant *in vivo* under high lipid supply and low energy demand. We conclude that, as long as lipid supply is low/modest, ACOT2 regulates matrix acyl-CoA levels in white SM to facilitate β-oxidation. When lipid supply is high, ACOT2 in white SM is still permissive for β-oxidation, but this is accompanied by the accretion β-oxidation intermediates and other lipids that can inhibit insulin signaling, though these effects might be lessened by higher proton leak.

### ACOT2 is not required for the mtFAS pathway

We could not find evidence that ACOT2 is required for the mtFAS pathway. The mtFAS pathway is well-known as the source of lipoic acid used as a co-factor by several matrix enzymes (Hiltunen, Chen et al., 2010). More recently, longer chain fatty acids were shown to also be a product of mtFAS (Nowinski, Solmonson et al., 2020), though the identity of the thioesterase that cleaves long-chain acyl groups from acyl carrier protein is unknown. We took the opportunity to investigate this in Acot2-/- mice, but could find no supportive evidence, such as lower ETC capacity in SM, which would have been predicted (Nowinski et al., 2020), or compensatory mitochondrial biogenesis.

### ACOT2 facilitates or hinders β-oxidation in white SM mitochondria, depending on lipid supply and ATP demand

Experiments in isolated mitochondria from white SM, using modest and high lipid supply and a range of simulated ATP demand, revealed the scope of possible impact of ACOT2 on β-oxidation. When PCarn supply was modest and simulated ATP demand was high, ACOT2 loss was associated with higher phosphorylating JO_2_. Furthermore, the CK clamp demonstrated that conductance for PCarn was higher in Acot2-/- mitochondria. These differences in Acot2-/- mitochondria occurred without obvious changes in the abundance of matrix enzymes. Thus, the most straightforward explanation is that flux through β-oxidation was greater in Acot2-/- mitochondria because long-chain acyl-CoA was not diverted through ACOT2. In other words, when Acot2 is present, long-chain acyl-CoA would be diverted through ACOT2 at the expense of β-oxidation. However, an apparent conflict needs to be addressed, between the latter concept and the finding that β-oxidation is higher in ACOT2 *overexpressing* hepatocytes (Moffat et al., 2014). In hepatocytes, ACOT2 overexpression led to higher β-oxidation, and this was largely attributed to higher proton leak (i.e., driven by fatty acids released by ACOT2) (Moffat et al., 2014). ACOT2 could also drive greater β-oxidation if fatty acids released by Acot2 are reactivated by acyl-CoA synthetase on the OMM then oxidized. Consistent with this possibility, we and others showed that most of the palmitate formed from PCarn oxidation (indicative of thioesterase activity) was exported from mitochondria (Gerber, Aronow et al., 2006, Seifert, Bezaire et al., 2008), and that a re-activation cycle was possible in isolated mitochondria (Seifert et al., 2008) (though how fatty acids are exported is unclear). Data from hepatocytes overexpressing Acot2 were also consistent with a re-activation cycle (Moffat et al., 2014). Finally, the conditions used here - PCarn (and malate) without added CoA – did not favor the re-activation of fatty acids. Thus, the greater PCarn-driven JO_2_ in isolated Acot2-/- mitochondria supplied with modest [PCarn] and high [ADP] is not inconsistent with a role for ACOT2 in promoting β-oxidation (Moffat et al., 2014).

In mitochondria from white SM, increasing the [PCarn] from 20 to 40 µM did not further increase phosphorylating JO_2_ in Loxp samples and resulted in lower JO_2_ in Acot2-/- samples. A CoA limitation might explain the results in Loxp but not Acot2-/- mitochondria. Rather, the observations in Acot2-/- mitochondria are consistent with a kinetic model of β-oxidation, based on β-oxidation in liver mitochondria, showing that, with increasing substrate supply, β-oxidation becomes vulnerable to product inhibition and also to feedback inhibition of 3-hydroxy-acyl-CoA dehydrogenase by elevated NAD^+^/NADH (Martines, van Eunen et al., 2017, van Eunen, Simons et al., 2013). Our data suggest that inhibition of β-oxidation worsens in white SM mitochondria when ACOT2 is absent.

### ACOT2 in muscle facilitates β-oxidation in vivo

The studies in isolated mitochondria showed that, when ACOT2 is absent, diverting more long-chain fatty acyl-CoA into the β-oxidation pathway can increase β-oxidation when ATP demand is high (i.e., when there is little back-pressure on the ETC) and lipid supply is not excessive. With high lipid supply, β-oxidation is associated with increased ACoAs build, suggesting β-oxidation overload, when ACOT2 is absent. To determine how ACOT2 impacts β-oxidation *in vivo*, we measured steady state levels of β-oxidation intermediates in white SM and undertook IC studies, allowing us to evaluate substrate metabolism under a normal range of lipid supply, β-oxidation flux and energy expenditure. Indirect calorimetry showed no evidence for higher β-oxidation; rather, glucose oxidation was higher (higher RQ, reflecting higher VCO_2_). Likewise, C2C12 myotubes acutely depleted of ACOT2 and incubated with glucose and palmitate (plus carnitine) showed higher pyruvate oxidation. Thus both longer-term and acute loss of Acot2 led to higher mitochondrial glucose oxidation, but never to higher β-oxidation, under a range of lipid supply.

In Acot2-/- mice fed regular chow or a 60% fat diet, higher RQ (reflecting higher VCO_2_) was evident mainly during the dark (active) phase of the day, when energy demand is fueled by a mix of glucose and other substrates. In SM under nutrient replete conditions, fatty acids are expected to be the main additional source of fuel. The relative contribution of fatty acids and glucose depends on diet composition, and fatty acyl-CoA-derived acetyl-CoA would compete with pyruvate-derived acetyl-CoA for entry into the TCA cycle. Our IC and isolated mitochondria data suggest that, in this competition scenario, loss of ACOT2 results in less contribution of β-oxidation-derived acetyl-CoA; in other words, ACOT2 would facilitate the access of fatty acyl-CoA-derived acetyl-CoA into the TCA cycle. During the rest phase of the day (for mice, the light phase), energy demand is largely fueled by fatty acids. Because demand is low at this time, β-oxidation capacity can be sufficient to match energy demand even if capacity is curtailed by ACOT2 loss; we propose that this occurred in Acot2-/- mice fed regular chow or fasted overnight, evidenced by the similar RQ (near 0.7) in Loxp and Acot2-/- mice (despite the accumulation of β-oxidation intermediates in Acot2-/- SM). With HFD (i.e., high lipid supply), we propose that β-oxidation capacity becomes vulnerable to inhibition that is mitigated by ACOT2 (evidenced by studies in isolated mitochondria) such that, when ACOT2 is absent, β-oxidation can no longer match even the low energy demand of the rest phase, and the contribution of pyruvate-derived acetyl-CoA to the TCA cycle rises.

The accumulation of β-oxidation intermediates in Acot2-/- SM and the poor correlation between some adjacent reactions of β-oxidation when lipid supply was low/modest (i.e., chow feeding and fasting), indicate that Acot2 is relevant during the rest phase of the day in chow-fed and fasted mice, and that ACOT2 mitigates the build-up of β-oxidation intermediates. That build-up was especially evident for C6- and C8-species is consistent with a kinetic model of mitochondrial β-oxidation flux showing that the medium-chain ketoacyl-CoA thiolase is most vulnerable to inhibition (Martines et al., 2017). It is notable that some build-up of intermediates and loss of correlation were evident for most chain lengths. This suggests that ACOT13 (similar substrate profile as ACOT2 (Wei, Kang et al., 2009)) or ACOT9 (specificity for C3- and C4-CoA (Tillander et al., 2014)) did not compensate in the SM of chow-fed or fasted mice. Also, ACOT9 and ACOT13 are CoA-inhibited (Bekeova et al., 2019, Tillander et al., 2014). While the build-up of β-oxidation intermediates in Acot2-/- Quad indicates that more CoA was sequestered in the β-oxidation pathway, the CoASH level was either unchanged (chow-fed Acot2-/- mice), greater (fasted Acot2-/- mice), or greater relative to ACoAs (HDF-fed Acot2-/- mice). We note that CoASH was measured in whole SM. Yet estimates of [CoASH] in mitochondria *vs*. cytosol (Naquet, Kerr et al., 2020), and consideration of relative compartment sizes (Larsen, Nielsen et al., 2012), put the CoASH amount substantially higher in mitochondria *vs*. cytosol, suggesting that CoASH in whole SM mainly reflects the mitochondria pool. Thus, while we cannot rule out that the CoASH pool available for β-oxidation was smaller in Acot2-/- SM, our data and the above arguments suggest that mitochondrial CoASH was not less when ACOT2 was absent.

In Quads from HFD Acot2-/- *vs*. Loxp mice, lower levels of β-oxidation intermediates and CoASH, and more instances of strong correlation between adjacent β-oxidation reactions were unexpected findings in light of the observations in chow-fed and fasted Acot2-/- mice. A different approach - measuring total CoA after hydrolysis of acyl-CoAs - in a different cohort of HFD-fed mice revealed lower levels of total CoAs in Acot2-/- Quad, consistent with the mass spectrometry data. Possibly, the lower CoASH in Quad mitochondria from HFD-fed Acot2-/- mice resulted in substantial activation of ACOT9 and ACOT13; this would be expected to facilitate β-oxidation or to increase the levels of other lipids, depending on the fate of the released fatty acids. Yet, VCO_2_ and RQ were higher in HFD-fed Acot2-/- mice during the light phase (and dark phase), consistent with higher glucose oxidation, not β-oxidation. Furthermore, lipidomics analysis revealed either no change (di- and triacylglycerides) or a decrease (ceramides, hexosylceramides and sphingomyelins) in lipids in Acot2-/- SM. Triacylglycerides were also unchanged in liver and heart of Acot2-/- mice *vs*. Loxp. Altogether, these observations indicate that, under high lipid supply, β-oxidation overload in white SM, *in vivo*, is mitigated by ACOT2 loss, though β-oxidation may be more susceptible to CoA limitation because CoASH was lower. Experiments to better understand how CoA is regulated in SM are warranted.

### Muscle ACOT2 contributes to proton leak in mitochondria when lipid supply is high

Experiments in mitochondria from white SM showed that Acot2 loss leads to lower proton leak, measured as lower non-phosphorylating JO_2_ at a given driving force (Δψm). This occurred when 40 µM PCarn was supplied, but not 20 µM PCarn. Likewise, there was evidence for higher proton leak when Acot2-overexpressing hepatocytes oxidized palmitoyl-CoA, and, *in vivo*, in fasted mice overexpressing Acot2 in hepatocytes (Moffat et al., 2014). Collectively, these data from loss- and gain-of-function models indicate that Acot2 can promote proton leak when lipid supply is high.

Evidence *in vivo* that endogenous muscle Acot2 contributes to proton leak in SM mitochondria is provided by the higher VO_2_ during the light phase in HFD-fed Loxp mice *vs*. Acot2-/- mice. That an effect of proton leak would be most evident when energy demand is low is consistent with the disproportionately higher proton leak contribution to JO_2_ at high Δψm (Nobes, Brown et al., 1990). Yet, an important caveat when considering proton leak *in vivo* is that substrate type and availability, and Δψm, cannot be accounted for. Nonetheless, that ACOT2 may be a source of fatty acids mediating proton leak is interesting because the ability of fatty acids to mediate proton leak across the IMM has long been known (Divakaruni & Brand, 2011), they can activate proton leak by the adenine nucleotide translocase (Bertholet, Natale et al., 2022)), and by uncoupling protein 1 in brown adipose tissue (Nicholls, 2021), but the source of the fatty acids is rarely considered. An ACOT2-mediated proton leak could have important implications; e.g., higher proton leak driven by ACOT2 could mitigate the build-up of β-oxidation intermediates, though the higher level of ACoAs in Quad from HFD *vs*. Chow-fed Loxp mice indicates that an ACOT2-mediated proton leak would not prevent ACoA build-up.

### Absence of ACOT2 in muscle improves systemic glucose homeostasis under HFD

Glucose homeostasis has been linked to mitochondrial β-oxidation in SM (Anderson et al., 2009, Finck et al., 2005, Gavin et al., 2018, Koves et al., 2008, Muoio & Neufer, 2012, Smith et al., 2021), and to cellular lipids (Petersen & Shulman, 2018). Because our findings revealed a role for endogenous Acot2 in β-oxidation in SM, it was of interest to evaluate glucose homeostasis and lipids SM. The chow-feeding and fasting conditions indicated that ACOT2 facilitates β-oxidation, and lower glucose oxidation. It might be tempting to speculate that preventing β-oxidation overload leads to improved glucose disposal. Yet, the ACoA build-up in Acot2-/- SM was not accompanied by any obvious defect in glucose disposal. On the other hand, under short term HFD, ACOT2 loss was protective against insulin resistance. Though the latter could be related to the lower ceramide levels in Acot2-/- SM, it should be noted that ceramides did not accumulate in SM from HFD-fed Loxp mice. Rather, hexosylceramide and sphingomyelin pools were expanded in Loxp SM, but not in Acot2-/- SM. Overall, these data suggest that Acot2-/- white SM is less at risk of ceramide accumulation when lipid supply is high. Finally, the mechanistic link between ACOT2 and sphingomyelins and ceramides is unclear. While ceramide and sphingolipid biosynthetic or hydrolyzing enzymes have been found in mitochondria of some cell types (Hernandez-Corbacho, Salama et al., 2017), we could not find evidence in SM mitochondria, despite finding several pathway members in heart mitochondria (unpublished). Thus, direct competition between ceramide or sphingolipid biosynthetic enzymes and ACOT2 in SM mitochondria seems unlikely. Even if indirect, our findings support as association between mitochondrial β-oxidation and the ceramide/sphingomyelin pools in white SM.

### Acyl/acetyl-CoA shunts and buffers

This study has revealed ACOT2 to be a *bona fide* regulator of matrix acyl-CoA concentrations via the interaction of Acot2 with β-oxidation. That this was especially evident in glycolytic SM, despite the relatively low protein expression of ACOT2 in glycolytic *vs*. oxidative SM might seem surprising. Yet, a recent study describing a shunt that mitigates acetyl-CoA build-up – by reversal of medium chain keto-acyl thiolase (MK, encoded by *Acaa2*) leading to C4-acyl-CoA generation determined that MK expression and the availability of this shunt were lower in glycolytic *vs*. oxidative muscle (Koves, Zhang et al., 2023). Similarly, expression of carnitine acetyl-transferase (CRAT), which also buffers C2- to C4-CoAs (Muoio, Noland et al., 2012), is lower in glycolytic *vs*. oxidative muscle (Koves et al., 2023, Murgia et al., 2015). Thus, altogether, these studies suggest that glycolytic SM may be especially sensitive to lipid overload in part because of the low expression of the individual shunt/buffer mechanism, leading to a low capacity for compensation in glycolytic SM. Another factor might be the increased expression of HADHB relative to ACADL in glycolytic *vs*. oxidation SM that we show in this study. We propose that, in glycolytic *vs*. oxidative muscle, each shunt and buffer is more important for mitigating lipid overload, Acot2 being one of them.

In conclusion, our data support a model in which ACOT2 facilitates β-oxidation in white SM when energy demand and lipid supply are low-to-modest. When lipid supply is high, ACOT2 does not prevent ACoA build-up (β-oxidation overload), and enables the accumulation of ceramide derivatives in white SM. Finally, the impact of ACOT2 loss in striated muscle on glucose homeostasis and SM lipids under high lipid supply raises the question of whether ACOT2 inhibition could be a strategy for mitigating some deleterious effects of lipid overload.

## MATERIALS and METHODS

### Mouse model

All experiments in mice were approved by the Institutional Animal Care and Use Committee of Thomas Jefferson University, protocol #01307. Mice were housed at 22°C, under a 12-hr light-dark cycle (lights on: 7:00 AM to 7:00 PM). Mice (C57BL/6, with intact Nnt gene) harboring floxed Acot2 alleles (exon 2 was floxed) (Bekeova et al., 2019) were crossed with mice expressing the striated muscle-specific CKM promoter to downregulate Acot2 expression in striated muscle (Acot2-/-). Experiments were performed on 11–14-week-old male mice. Different groups of mice were used as: ad libitum fed a standard diet (LabDiet 5001, Purina), mice that were fasted overnight (5:00 PM to 9:00 AM) or mice fed a HFD (Open Source Diet, D12492, Rodent Diet with 60 kcal% Fat) for 7 days before harvesting the tissue or isolating mitochondria. Mitochondria were isolated or whole quadriceps samples were harvested always in the morning, either used for bioenergetics studies or snap frozen, and stored at -80°C.

### Glucose tolerance test

The chow fed mice were fasted overnight, blood glucose level was measured before the oral injection of 2g of glucose for 1kg of body weight. The HFD mice were fasted for 6h, blood glucose level was measured before the oral injection of 1 g of glucose for 1 kg of body weight. Glucose clearance was measured at 15, 30, 60, 90, 120 min after glucose delivery. The blood sample was obtained from a small nick at the end of the mouse’s tail. Glucose level was analyzed using a glucose meter (Abbot).

### Insulin tolerance test

Chow fed mice were fasted overnight; blood glucose level was measured before the IP injection of 1.5U/kg insulin. The HFD mice were fasted for 6h; blood glucose level was measured before the IP injection of 3U/kg of insulin. In both chow- and HFD-fed mice, blood glucose level was also measured in 10 min after insulin injection.

### Treadmill running

Mice were run on treadmill (Columbus Instruments) before the collection of quadriceps muscle for acyl-CoA analysis. The protocol was the following: 20 min, 10 m/min, 10% incline.

### Indirect calorimetry

Indirect calorimetry studies were performed at the University of Pennsylvania Diabetes Research Center Rodent Metabolic Phenotyping Core using the Promethion system (Sable Systems, Las Vegas, NV). Each cage is equipped with an XYZ beam break array (BXYZ-R, Sable Systems), and mass measurement modules (2 mg resolution) for food intake and water intake. The set-up was housed in temperature-controlled unit (20-22°C), maintained on a 12h light-dark cycle. Mice were acclimated for 5 days before recording. Oxygen consumption (VO_2_) and carbon dioxide (VCO_2_) production were measured for each mouse at 5 min intervals for 20 seconds resulting in a 3 min cycle time. Respiratory quotient is calculated as the ratio of VCO_2_/VO_2_. Data acquisition and instrument control were controled by the IM-3 software, and the raw data were processed using MacroInterpreter v.2.38 (Sable Systems).

### Isolation of skeletal muscle mitochondria

Skeletal muscle isolation (Vasquez-Trincado, Dunn et al., 2022) was performed on ice or at 4°C. After the skeletal muscle was dissected and washed in isolation buffer (IB: 140 mM KCl, 20 mM HEPES, 5 mM MgCl2, 1 mM EGTA, pH 7), the tissue was minced with a razor blade. This was followed by resuspension in IB + 1 mM EGTA, 1 mM ATP, 1% w/v defatted BSA (Sigma, A7030) in a glass/Teflon Potter-Elvehjem homogenizer; and homogenization (500 rpm, 16 passes). Samples were centrifuged 10 min, 2000 rpm (478g) (Sorvall SS-34 rotor), in the next step the supernatant was centrifuged 10 min, 9000 rpm (9681g), then the pellet was resuspended in 1 ml of IB, with addition of 24 ml of IB the sample was incubated on ice for 5 min. In the next step the sample was centrifuged 10 min, 2000 rpm, the supernatant was filtered through 70 um nylon cell strainer and centrifuged 10 min, 9000 rpm. The final pellet was resuspended in IB in a volume that resulted in a protein concentration of ∼25 mg/ml. The protein concentration was determined using bicinchoninic acid assay (BCA).

### Bioenergetics analyses in isolated SM mitochondria

O2 consumption (JO2) was measured using the Seahorse XFe24 Analyzer (Seahorse Bioscience, Billerica, MA, USA)(Moffat et al., 2014, Vasquez-Trincado et al., 2022). Immediately after the isolation, fresh mitochondria were used for the measurement. Each well of the custom microplate contained 5 μg of mitochondria for the condition with pyruvate as substrate and 7 μg of mitochondria for the condition with palmitoyl-L-carnitine as substrate. Mitochondria were suspend in mitochondria assay medium (MAS: 70 mM sucrose, 22 mM mannitol, 10 mM KH2PO4, 2 mM Hepes, 1 mM EGTA, 5 mM MgCl2, 0.2% defatted BSA, pH 7.4 at 37°C) containing the appropriate substrate. Amount of mitochondria had been optimized such that the O2 *vs*. time signal was linear under all conditions. To promote adhesion of mitochondria to the plastic the microplate was centrifuged at 2000*g*, 20 min, 4°C. The attachment was verified after centrifugation and again after experiments. During the experiment different substrates were tested: pyruvate/malate (10mM/10 mM), palmitoyl-L-carnitine/malate (20 μM/0.1 mM; 40 μM/0.1 mM; 40 μM/0.3 mM; 40 μM/3 mM; 30 μM/0.2 mM; 30 μM/10 mM). The following injections were used: ADP 2.8 mM/well (state 3), oligomycin (4 µg/ml) was used to measure non-phosphorylation “leak” respiration JO2 (state 4) and the uncoupler FCCP (4 μM) was used to measure maximal electron transport chain activity.

### Oroboros – CK clamp

We used an adopted method from Fisher-Wellman et al., 2018 for our measurements. Baseline JO2 was recorded with mitochondria, 200 µg of isolated mitochondria in 2ml of reaction volume, in MAS buffer (MAS: 70 mM sucrose, 22 mM mannitol, 10 mM KH2PO4, 2 mM Hepes, 1 mM EGTA, 5 mM MgCl_2_, 0.2% defatted BSA, pH 7.4 at 37°C) supplemented with 5 mM ATP, 5 mM creatine, 1 mM phosphocreatine, 20 U/ml creatine kinase. Followed by substrate addition, 0.01 M pyruvate and 0.01M malate, or 0.04 mM Pcarn and 0.3 mM malate to stimulate JO2. Then we added, as many times as needed, 3 mM phosphocreatine (3×3 mM PCr and 5×10 mM PCr for PcCarn/Mal as a substrate) at a time to decrease JO_2_ back towards the baseline.

### Thioesterase activity assay

Samples were diluted to 20 μg/μl in mitochondrial isolation buffer and subjected to five freeze-thaw cycles. Reactions were run in a total volume of 300 μl in reaction buffer (50 mM KCl, 10 mM HEPES pH 7.8) at 37°C. To prewarmed buffer 40 μg of the sample/well was added and 0.3 mM DTNB. Absorbance was measured at 412 nm every 1 s for 3 min. Then substrate was added and absorbance was measured at 412 nm every 1 s for 5 min.

### Immunoblot analysis

Western blot analysis were done on isolated mitochondria samples or tissue samples quickly frozen in liquid nitrogen. The quadriceps tissue was homogenized with a lysis buffer containing: NaCl, 150 mM; HEPES 25 mM; EGTA 2.5 mM; Triton 100X 1%; Igepal 1%; SDS 0.10%;

Sodium deoxycholate 0.50%; Glycerol 10%, Protease inhibitor (Roche 11873580001) and Phosphatase inhibitor cocktail (sodium fluoride 200mM, imidazole 200mM, sodium molybedate 115mM, sodium orthovanadate 200mM, sodium tartrate dihydrate 400mM, sodium pyrophosphate 100mM and β-glycerophosphate 100mM), using a glass/Teflon homogenizer at 300 rpm, followed by incubation on ice for 45 min, then spin at 18000g at 4°C for 20 min. From the supernatant, protein concentration was measured by bicinchoninic acid assay (BCA). Samples were loaded onto a 10% or 12% polyacrylamide gel (for Licor-conjugated secondary antibodies) or a 4-12% Bis-Tris gel (chemiluminescence (ECL)-conjugated secondaries), electrophoresed, transferred onto nitrocellulose membrane and blocked with Odyssey blocking buffer diluted in PBS (LI-COR Biosciences, Lincoln, NE, USA) or with TBS-T plus 3% BSA. The membrane was incubated overnight in primary antibodies (see Table, below) diluted in TBS-T with 0.02% NaN3. Secondary antibodies were diluted 1:20000 in 5% skim milk (LI-COR anti-MS 92632212 and anti-Rb 92632213) or in TBS-T 1:50000 for ECL (Thermo Fisher anti-MS 31473 and anti-Rb 31460). Protein bands were visualized using the Azure Biosystem c600.

Primary Antibodies used in immunoblot analysis

| Antibody | Catalog Number | Dilution | Host |
| --- | --- | --- | --- |
| ACOT2 | Custom Antibody (see (Bekeova et al., 2019)) | 1:1000 | Rabbit |
| ACOT9 | Custom Antibody (see (Bekeova et al., 2019)) | 1:1000 | Rabbit |
| ACOT13 | Custom Antibody (see (Bekeova et al., 2019)) | 1:1000 | Rabbit |
| VLCAD | Ab128566 | 1:1000 | Mouse |
| β-HAD | Ab203114 | 1:1000 | Rabbit |
| ETFa | Ab153722 | 1:1000 | Rabbit |
| NUDT8 | Custom Antibody (see (Kerr, Shumar et al., 2019)) | 1:1000 | Rabbit |
| COASY | Custom Antibody (see (Kerr et al., 2019)) | 1:1000 | Rabbit |
| Phospho-PDH (Ser293) | Novus NB110-93479 | 1:1000 | Rabbit |
| PDH | Abcam 110330 | 1:1000 | Mouse |
| Phospho-ACC (Ser79) | CST-3661L | 1:1000 | Rabbit |
| ACC | CST-3676T | 1:1000 | Rabbit |
| GRB2 | SC-8034 | 1:1000 | Rabbit |

### ELISA - Insulin measurement

The chow fed mice were fasted overnight, the HFD mice were fasted for 6h before the blood samples were collected for serum isolation. Samples were obtained from a small nick at the end of the mouse’s tail. They were incubated on RT for 30min, centrifuged on 4°C, 1500g, for 10min and the blood serum was transferred into clean tubes, stored on -80°C until the experiment. Samples were also collected after 15 min of the oral injection of 2ug of glucose for 1g of body weight of chow fed mice and 1ug of glucose for 1g of body weight of HFD mice. For the insulin measurement we used ALPCO Insulin Rodent Chemiluminescence ELISA kit. The measurement and the analysis were done based on the protocol included in the kit.

### CoA extraction, derivatization and quantification by HPLC

The samples were processed as previously described in (Zano, Pate et al., 2015). For the total CoA measurements we used 30-50 mg of quadriceps tissue. The samples were homogenized in 2 ml of ice cold 1 mM KOH for 30 seconds at 600 rpm in a glass/Teflon Potter-Elvehjem homogenizer. This step was followed by an addition of 500 μl of 125 mM KOH to bring the pH ≈ 12 and vortex. The tubes with the samples were parafilmed and incubate at 55°C for 2 h, to hydrolyze CoA thioesters to free CoA. In the next step we added 160 μl of 1M Trizma-HCl pH∼ 3.9 to bring the pH ≈ 8, which is optimal for the reaction with mBBr. We added 10 μl of 100 mM mBBr (Echelon Biosc. F-0030) dissolved in acetonitrile and incubated the samples at RT in dark for 2h. In next step we acidified the samples with the addition of 100 μl of glacial acetic acid, centrifuged the samples at 500 g for 10 minutes at 27°C. The supernatant was applied on an already equilibrated 2-(2-pyridyl)ethyl columns (Sigma 54127-U) with 1 ml of 50% MeOH with 2% acetic acid. In the next step the columns were washed with 2 × 1 ml of 50% MeOH with 2% acetic acid and 1 ml of water. The fraction containing derivatized CoA was eluted with 2 × 1 ml of 95% ethanol with 50 mM ammonium formate and then 2 × 1 ml of 90% methanol with 15 mM ammonium hydroxide into a glass vial. Samples were dried under nitrogen gas, resuspend in 300 μl of water by vortexing and incubating at room temperature for 10 minutes. In next step the samples were transferred to a Spin-X Centrifuge Tube Filter (Costar, Cat# 8161, 0.22 µm Cellulose Acetate) and centrifuged at 3000 g for 10 minutes. At the end the samples were transferred to HPLC vial. We prepared mBBr-CoA standard. 2 mM Free CoA (Sigma C3019) was combined with 2 mM mBBr in 100 mM TrisHCl pH = 8 and incubated in dark at RT for 2 h. The resulting mBBr-CoA was used for constructing the standard curve. The samples were analyzed by HPLC, as described by Shumar et al., 2015, using a Waters Alliance e2695 system equipped with a 2489 UV/Vis detector and a Kintex 4.6 × 150 mm, 3 μm C-18 column (Phenomenex) kept at 35°C. Fifteen microliters of the samples were injected onto the column, and the flow rate was 0.5 ml/min. The HPLC elution was the following: 0–2.0 min, 90% buffer A (50 mM KH2PO4, pH4.6), 10% buffer B (acetonitrile); 2.0–9.0 min concave gradient to 25% B; 9.0–23 min linear gradient to 40% B; 23.0–30 min linear gradient to 90% A and column equilibration in 90% A. Monobromobimane-CoA was detected by monitoring the absorbance at 393 nm and quantified using a standard curve.

### Analysis of acyl-CoAs (absolute values) of β-oxidation intermediates

Acyl-CoAs were extracted by the method of Minkler, *et al*., (Minkler, Kerner et al., 2008) using internal standards generated as previously described (PMID: 25572876) and spiked into each sample and calibration standard, then analyzed by liquid chromatography-high resolution mass spectrometry (LC-HRMS) using the method validated previously (Frey, Feldman et al., 2016, Snyder, Basu et al., 2014). Briefly, previously harvested heart samples were weighed still frozen and 650 µl of acetonitrile:isopropanol (3:1 v/v) and 100 µl of the yeast acyl-CoA internal standard library (derived from pan6-deficient *Sacharomyces cerevisiae* grown in [^13^C_315_N_1_]-pantothenic acid-containing media) were added to the samples. Samples were vortex mixed then probe tip sonication, five times 30 pulses with ½ second delay, followed by 5 min vortexing, and 10 min sonication in water bath to homogenize the tissue. Next 250 µl of 100 mM KH_2_PO_4_ (pH 6.7, prepared fresh) was added to the samples, vortex mixed, then centrifuged at 16000 × *g* for 5 min at 4°C. The supernatant was transferred into a glass tube and 125 µl of glacial acetic acid was added to acidify the samples before solid phase extraction (SPE). 2-(2-pyridyl)ethyl-functionalized silica SPE columns were conditioned with 1 ml of acetonitrile: isopropanol: water: acetic acid (9:3:4:4 v/v/v/v), the acidified samples were loaded on the columns, the columns were washed with 2 ml of acetonitrile:isopropanol:water:acetic acid (9:3:4:4 v/v/v/v), then eluted into fresh glass tubes with 1 mL methanol:250 mM ammonium formate (4:1 v/v). The samples were evaporated to dryness under nitrogen gas and re-dissolved in 100 μl of water:acetonitrile (7:3 v/v) for LC-HRMS analysis.

5 µL injections of the extracted acyl-CoAs were analyzed on an Ultimate 3000 Quaternary UHPLC coupled to a Q Exactive Plus mass spectrometer operating in the positive ion mode with a heated ESI probe in an IonMax Source housing as previously described (Frey et al., 2016). Samples were kept in a temperature controlled autosampler at 6 °C and LC separation was performed as previously described on a Waters XBridge 3.5 μm particle size C18 2.1 × 150 mm column. LC conditions were as follows modified from previous studies (Snyder et al., 2014); column oven temperature 25 °C, solvent A water with 5 mM ammonium acetate, solvent B 95:5 acetonitrile: water with 5 mM ammonium acetate, solvent C (wash solvent) 80:20 acetonitrile: water with 0.1% formic acid. The gradient was as follows: 0.2 mL/min flow at 98% A and 2% B for 1.5 min, 80% A 20% B at 5 min, 100% B at 12 min, 0.3 mL/min 100% B at 16 min, 0.2 mL/min 100% C at 17 min, held to 21 min, then re-equilibrated at 0.2 mL/min flow at 98% A and 2% B from 22 to 28 min. Flow from 4–18 minutes was diverted to the instrument. Operating conditions on the mass spectrometer were as follows; auxiliary gas 10 arbitrary units (arb), sheath gas 35 arb, sweep gas 2 arb, spray voltage 4.5 kV, capillary temperature 425 °C, S-lens RF-level 60, aux gas heater temperature 400 °C, in-source CID 5 eV. Acquisition used alternating full scans from 760–1800 m/z at 140,000 resolution and data independent acquisition looped 3 times with all fragment ions multiplexed at a normalized collision energy of 20 at a resolution of 280,000.

### Organic acids and amino acids extraction and LC-MS

Quadriceps tissue samples were harvested and quickly frozen in liquid nitrogen. The samples were powdered on dry ice and ∼25 mg of the powder was used for the extraction. We used the protocol from Chicco et al., 2018 with some modification. To each sample was added 1 ml of MeOH:ACN:water (5:3:2 v/v/v) and incubated on ice for 30 min with vortexing every 5 min. Then the samples were centrifuged at 10000xg and 4 °C for 15 min, and the supernatant was transferred to clean Eppendorf tubes. Samples used to detect citric acid, glutamine and serine were directly used for LC-MS analysis. Samples used for the detection of leucine, isoleucine, valine and glycine were further concentrated and then used for analysis. Samples for detection of aKG were concentrated and derivatized. Samples for detection of succinic acid and malic acid were directly derivatized after extraction. The concentration step was combined with deproteinization. The samples were mixed with ACN (1:3 v/v), vortexed for 1 min, incubated at - 20 °C for 20 min then dried overnight in a hood. The samples were resuspended in 1/10 of the original extract volume. Before injection, the samples were centrifuged for 30 min at 4 °C and 10000g. In the derivatization step (adopted from Yip et al., 2017) 30 µl of the extract was used. 2 µl of 1 M aniline (in 1:1 v/v water/ACN) was added (pH > 7) 2 µl of 1 M EDC (dissolved in water) was added. The samples were incubated overnight at 4 °C, then quenched with 1 µl of formic acid (pH < 7). Before injection the samples were centrifuged for 30 min at 4 °C and 10000xg. Metabolites were analyzed on a Waters Acquity I-Class UPLC system coupled to a Synapt G2Si HDMS mass spectrometer in positive ion mode with a heated electrospray ionization (ESI) source in a Z-spray configuration. LC separation was performed on a Waters Acquity UPLC BEH 1.7 µm 2.1 × 50 mm column equipped with a Vanguard C18 precolumn. For derivatized metabolites a 0.6 mL/min gradient of 95/5 to 15/85 A/B in 4 min followed by washing and reconditioning the column, whereas for polar metabolites a 0.6 mL/min isocratic run at 95/5 A/B was used. Eluent A is 0.1% v/v formic acid in water and B is 0.1% v/v formic acid in ACN. Conditions on the mass spectrometer were as follows: capillary voltage 0.5 kV, sampling cone 40 V, source offset 80 V, source 120 °C, desolvation 250 °C, cone gas 0 L/h, desolvation gas 1000 L/h and nebulizer 6.5 bar. The analyzer was operated in resolution mode and low energy data was collected between 100 and 1500 Da at 0.2 sec scan time. MS^e^ data was collected using a ramp trap collision energy 20-40 V, and masses were extracted from the TOF MS TICs using an abs width of 0.005 Da. Peak picking and integration was automated using Waters Unifi 1.4.

### Lipidomics analysis

Optima grade methanol, water, acetonitrile, methyl tert-butyl ether and 2-propanol were from Thermo Fisher Scientific (Pittsburg, PA). Gasses were supplied by Airgas (Philadelphia, PA). Glassware and HPLC vials were from Waters Corp (Milford, MA).

#### Tissue preparation for lipids extraction

Pieces of 20-30 mg of frozen tissues were cut on a tile kept in dry ice with a new blade kept in dry ice. The tissue was added to low retention Eppendorf tube prepared with 0.6 mL 80% methanol (MeOH) and 20 µL SPLASH® LIPIDOMIX (diluted 1:1 with methanol from original #330707 Avanti solution) (Avanti Polar Lipids, Alablaster, AL) and kept in dry ice. Samples were pulse sonicated for 30x half-second pulse on ice and kept on ice for 20 min for metabolites extraction. Each tube was then vortexed 3x 30 seconds each. The tissue homogenates was then moved to a 10 mL glass Pyrex tube with screw cap. The Eppendorf tubes were rinse with 0.5 mL methanol and added to same glass tube. 5 mL methyl tert-butyl ether (MTBE) was added to each of the tubes and then tubes were shaken vigorously for 30 min. 1.2 mL water was added to each tubes and vortexed for 30 sec each. Centrifugation for 10 min @ 1000xg created two phases. The top clear phase was moved to a clean glass Pyrex tube and dried down under nitrogen. 100 µL MTBE/MeOH=1/3 (v/v) was used to re-suspend the residue. The sample was spun down at 10, 000 × g for 10 min at 4°C and only the top 50 µL were transferred to a HPLC vial for LC-MS analysis. A pooled sample was created by mixing 10 µL of each re-suspended sample. 2 µL injections were made.

#### Liquid chromatography high resolution -mass spectrometry (LC-HRMS) for lipids

Separations were conducted on an Ultimate 3000 (Thermo Fisher Scientific) using an Ascentis Express C18, 2.1 × 150 mm 2.7μm column (Sigma-Aldrich, St. Louis, MO). The flow-rate was 0.4 m /min, solvent A was water:acetonitrile (4:6 v/v) with 0.1% formic acid and 10 mM ammonium formate and solvent B was acetonitrile:isopropanol (1:9 v/v) with 0.1% formic acid and 10 mM ammonium formate. The gradient was as follows: 10 % B at 0 min, 10 % B at 1 min, 40 % B at 4 min, 75 % B at 12 min, 99 % B at 21 min, 99 % B at 24 min, 10 % B at 24.5 min, 10 % at 30 min. Separations were performed at 55 °C. For the HRMS analysis, a recently calibrated QE Exactive-HF mass spectrometer (Thermo Fisher Scientific) was used in positive ion mode with an HESI source. The operating conditions were: spray voltage at 3.5 kV; capillary temperature at 285°C; auxiliary temperature 370°C; tube lens 45. Nitrogen was used as the sheath gas at 45 units, the auxiliary gas at 10 units and sweep gas was 2 units. Same MS conditions were used in negative ionization mode, but with a spray voltage at 3.2 kV. Control extraction blanks were made in the same way using just the solvents instead of the tissue homogenate. The control blanks were used for the exclusion list with a threshold feature intensity set at 1e10^^5^. Untargeted analysis and targeted peak integration was conducted using LipidsSearch 4.2 (Thermo Fisher Scientific) as described by Wang et al (DOI: 10.4155/bio-2021-0098). An external mass calibration was performed using the standard calibration mixture approximately every three days. All samples were analyzed in a randomized order in full scan MS that alternated with MS2 of top 20, with HCD scans at 30, 45 or 60 eV. Full scan resolution was set to 120,000 in the scan range between *m/z* 250–1800. The pool sample was run every 15 samples. Lipids quantification was done from the full scan data. The areas were normalized based on the amount of the internal standard added for each class (see Table, below). All amounts were normalized to the original tissue weight.

| 330707 \| SPLASH® LIPIDOMIX® Mass Spec Standard |  |  |  |  |  |  |  |
| --- | --- | --- | --- | --- | --- | --- | --- |
|  | Splash LipidoMix Internal Standard | Formula | pmole/sample | m/z Positive | Adduct | Stock conc | Mol weight |
| IS-01 | 18:1(d7) Chol Ester | C45H71D7O2 | 5322 | 675.6777 | M+NH4 | 350 ug/mL | 657.65 |
| IS-02 | 15:0-18:1(d7) PC | C41H73D7NO8P | 2126 | 753.613 | M+H | 160 ug/mL | 752.61 |
| IS-03 | Cholesterol (d7) | C27H39D7O | 2542 | 376.3955 | M-H2O+H | 100 ug/mL | 393.41 |
| IS-04 | 15:0-18:1(d7)-15:0 TG | C51H89D7O6 | 678 | 829.7979 | M+NH4 | 55 ug/mL | 811.77 |
| IS-05 | 15:0-18:1(d7) PG | C39H67D7O10P | 405 | 758.5798 | M+NH4 | 30 ug/mL | 741.55 |
| IS-06 | 18:1(d9) SM | C41H72D9N2O6P | 407 | 738.6464 | M+H | 30 ug/mL | 737.65 |
| IS-07 | 18:1(d7) LPC | C26H45D7NO7P | 473 | 529.3991 | M+H | 25 ug/mL | 528.40 |
| IS-08 | 15:0-18:1(d7) DG | C36H61D7O5 | 170 | 605.5844 | M+NH4 | 10 ug/mL | 587.55 |
| IS-09 | 15:0-18:1(d7) PI | C42H75D7NO13P | 118 | 847.6034 | M+H | 10 ug/mL | 847.60 |
| IS-10 | 15:0-18:1(d7) PA | C36H61D7O8P | 105 | 685.5507 | M+H | 7 ug/mL | 667.52 |
| IS-11 | 15:0-18:1(d7) PE | C38H67D7NO8P | 70 | 711.5663 | M+H | 5 ug/mL | 710.57 |
| IS-12 | 15:0-18:1(d7) PS | C39H66D7NO10P | 64 | 755.5562 | M+H | 5 ug/mL | 777.54 |
| IS-13 | 18:1(d7) LPE | C23H39D7NO7P | 103 | 487.3524 | M+H | 5 ug/mL | 486.35 |
| IS-14 | 18:1(d7) MG | C21H33D7O4 | 55 | 364.3439 | M+NH4 | 2 ug/mL | 363.34 |

### Triglyceride extraction

Triglycerides were extracted from previously collected tissues (quadriceps, liver, heart), frozen in liquid nitrogen. Around 100 mg of the frozen tissue was used for the extraction. The samples were digested in 6x of the tissue weight of 100% EtOH: 30% KOH (2:1) solution, overnight in water bath at 60°C. The samples were vortexed before incubation. Next day 1.08x of the volume of the digestion solution of 1M MgCl_2_ was added to the samples. Which was flowed by vortexing and 10min incubation on ice. The samples were centrifuged at RT for 30 min at 14000g. The supernatant was diluted 5x and used for the measurement with Stanbio TG kit (2200-430 Triglyceride LiquiColor Mono).

### RNA sequencing

Quadriceps tissue samples were harvested and quickly frozen in liquid nitrogen. Samples were powered on dry ice and used for RNA isolation using Norgen Animal Tissue RNA Purification Kit (cat.25700).

*Library preparation, total RNA sequencing of chow fed and fasted samples:* Total RNA quality was assessed using an Agilent TapeStation (Agilent, Palo Alto, CA, USA) and RNA concentration was quantified by Qubit 4.0 spectrophotometer (Invitrogen, Carlsbad, CA, USA). The library was prepared for sequencing from 300 ng RNA of each sample using the Illumina Truseq standard total RNA library preparation kit (Illumina, San Diego, CA, USA). The quantity and quality of amplified libraries were evaluated using Qubit 4.0 and Agilent TapeStation high sensitivity D1000 Screen Tape. The average fragment length of the cDNA libraries was 330 bp. RNA-seq libraries were sequenced using paired-end (2×75) and high output flow cell v2.5 on Illumina NexSeq 500 instruments following the manufacturer’s protocol.

#### Library preparation, total RNA sequencing of HFD samples

*We* sent the samples to Genewiz/ Azenta for Standard RNA-seq analysis.

*To analyze all the raw data* we used https://www.rosalind.bio/.

### Proteomics

The mass spectrometry-based proteomics data that were used to generate Figure 1A can be found in Supplementary Tables S1 and S2 of Murgia et al (Murgia et al., 2015). The corresponding raw and processed files are available for downloading from the PRIDE Proteomics Identification Database with the dataset identifier PXD001641.

### Bioenergetics analyses in C2C12 myotubes

O_2_ consumption (JO_2_) was measured using the Seahorse XFe24 Analyzer (Seahorse Bioscience, Billerica, MA, USA). C2C12 cells were cultured in DMEM media with 4.5g/l D-glucose, L-glutamine, 110 mg/l sodium pyruvate (Gibco 11995-065) with addition of 10% FBS and 1% Penicillin/Streptomycin. The differentiation was initiated with the change of media containing 2% FBS. The seahorse measurement was ran on the seventh day of differentiation. We used knockdown cells, which were prepared using 10 pmol/well (of 6 well cell culture plate) Acot2 or control siRNA (Origene) and 1.5ul of Lipofectamine RNAiMAX using forward transfection. For part of the experiments the cells were overnight cultured in substrate limited media (DMEM Sigma D5030, 3.7 g/l NaHCO_3_, 0.5 mM glucose, 1 mM glutamine, 0.5 mM carnitine, 1% FBS pH 7.4 at 37°C). As an assay buffer we used a media optimized for measuring β-oxidation in C2C12 cells (111 mM NaCl, 4.7 mM KCl, 1.25 mM CaCl_2_, 2 mM MgSO_4_, 1.2 mM NaH_2_PO_4_) with additional supplements the day of the assay (0.5 mM carnitine, 5 mM HEPES, required substrates: 10 mM glucose, 2 mM glutamine, 100 µM palmitate conjugated to BSA (5 mM Palmitate: 0.8 mM BSA (6:1 palmitate:BSA) in 150 mM sodium chloride, pH 7.4)) pH 7.4 on 37°C. Carnitine was not used when just glucose was present as substrate. A variety of injections were used: oligomycin (0.465 ug/ml /well) was used to measure non-phosphorylation “leak” respiration JO2, the uncoupler FCCP (1uM/well) was used to measure maximal electron transport chain activity, 2-DG (50mM/well) was used to confirm that the ECAR produced during the experiment was due to glycolysis, Etomoxir (3uM/well) was used to inhibit CPT1 and measure OCR by other substrates then long chain FA (palmitoyl we added), UK5099 (2uM/well) was used to inhibit MPC to inhibit pyruvate dependent O2 consumption.

### Statistical analysis

Data are presented as mean ± standard error of the mean, and as individual points from each mouse or cell. Statistical analysis was performed in GraphPad Prism 9 (GraphPad Software, San Diego, CA). Statistical significance between 2 groups was determined using 2-tailed unpaired t-tests. For experiments with a 2×2 design (i.e., genotype x condition), 2-way ANOVA was using, together with pairwise comparisons using the Benjamini, Krieger, Yekutieli method to correct for multiple comparisons, using a false discovery rate of 0.05. Further details of statistical testing and p or q values are provided in the figure legends.

## ACKNOWLEDGEMENTS

Funding for this project: NIH R01 DK109100 (ELS), NIH R01 GM132261 (NWS), NIH R35 GM119528 (RL), NIH R01 AA018873 (RV), NIH P30 ES013508 (CM).

## Author Approvals

All Authors have seen and have approved the manuscript, and the manuscript has not been accepted or published elsewhere.

