## Supplemental Figures for "Acyl-CoA thioesterase-2 facilitates β-oxidation in glycolytic skeletal muscle in a lipid supply dependent manner"

### Supplemental Figure 1

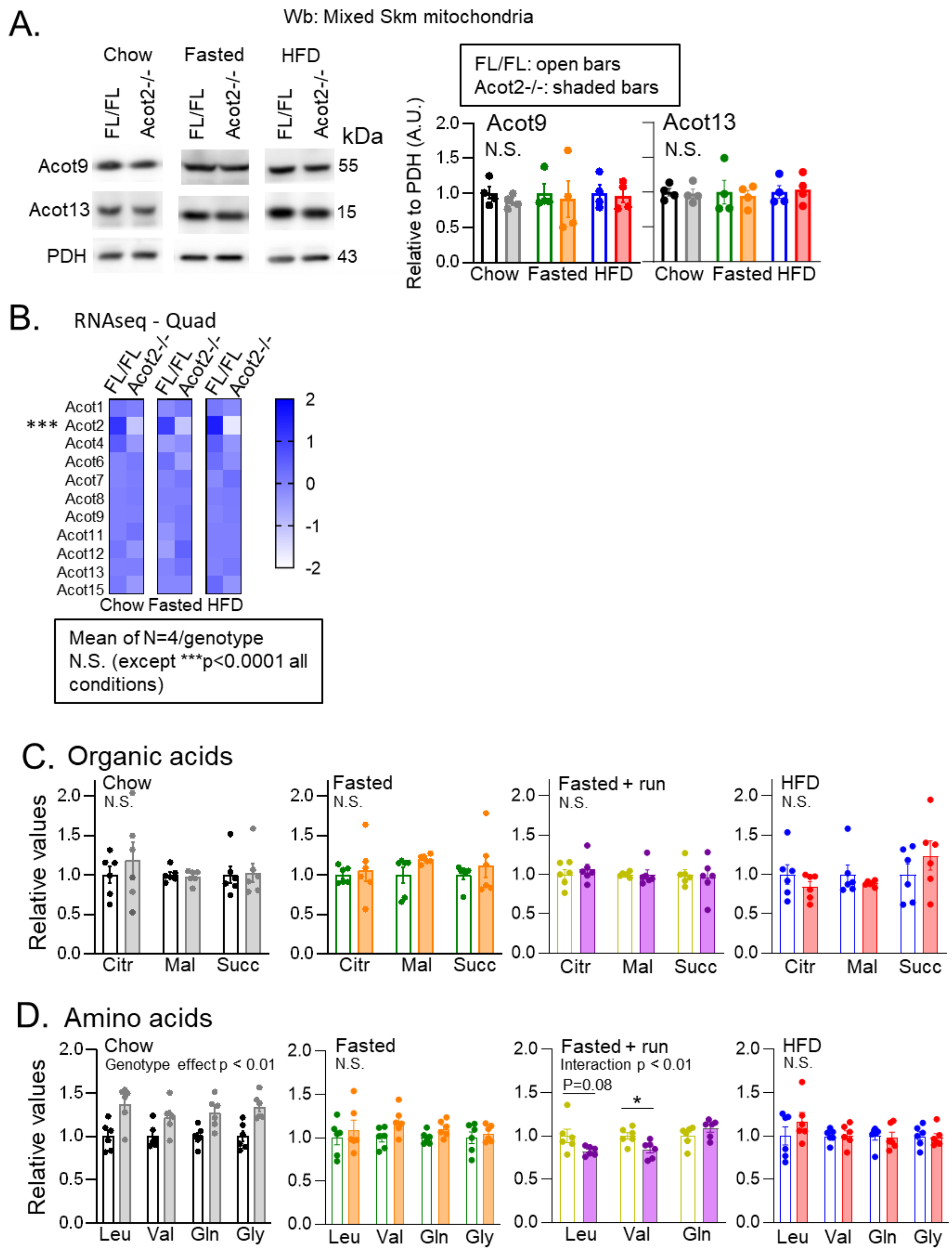

**Suppl. Fig. 1. Lack of change in expression of other Acots in Quad from mice with Acot2 loss in striated muscle**

A. Representative immunoblots for Acot9 and Acot13 in FL/FL and Acot2-/- SKM mitochondria under chow, fasted and HFD conditions, and quantification in n=4/genotype with PDH as loading control. N.S. not significant (two-way ANOVA).

B. Transcript levels of each Acot2, from RNAseq analysis. RNA was isolated from Quad from mice fed standard chow, overnight fasted, of fed HFD for 7 days.

C, D. Mass spectrometry of quadriceps samples collected from FL/FL (wild type) and Acot2-/- (Acot2 knockdown) mice studied under chow, overnight fasted, run after overnight fasting and 7-day HFD conditions in n=6/genotype. Comparison of the relative abundance of selected amino acids (C), and organic acids (D). Statistics: two-way ANOVA (p values) and multiple comparisons (* q=0.04) with correction for FDR (0.05; Benjamini, Krieger, Yekutieli method). N.S. not significant.

Panel A, C, D: Bars are mean ± s.e.m, the points on the bar graphs represent individual mice.

### Supplemental Figure 2

A. Total CoAs, derived from all CoA species - Quad

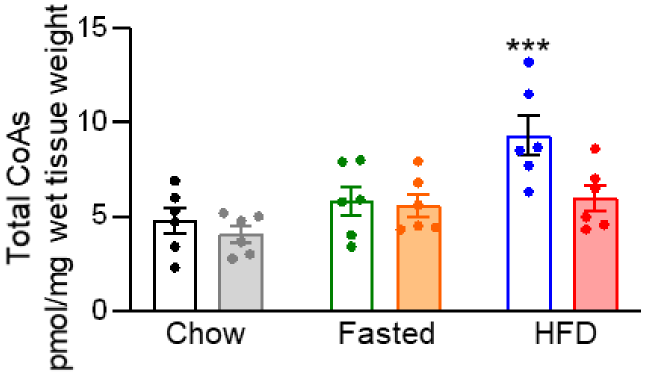


Interaction p = 0.09
Genotype effect p = 0.02
Diet effect p = 0.0005
***: q < 0.05 *vs.* all other conditions (FL/FL and Acot2-/-)

FL/FL: open bars
Acot2-/-: shaded bars

**Suppl. Fig. 2. HFD feeding lowers the abundance of total CoAs in Acot2 depleted Quad**

Total CoA abundance was measured by HPLC, as described in Methods, in Quads from mice fed standard Chow, overnight fasted, or fed HFD for 7 days. Statistics: two-way ANOVA and multiple comparisons (q values shown in the Figure) with correction for FDR (0.05; Benjamini, Krieger, Yekutieli method). N=6/genotype/condition.

### Supplemental Figure 3

A. Chow FL/FL vs. Acot2-/-

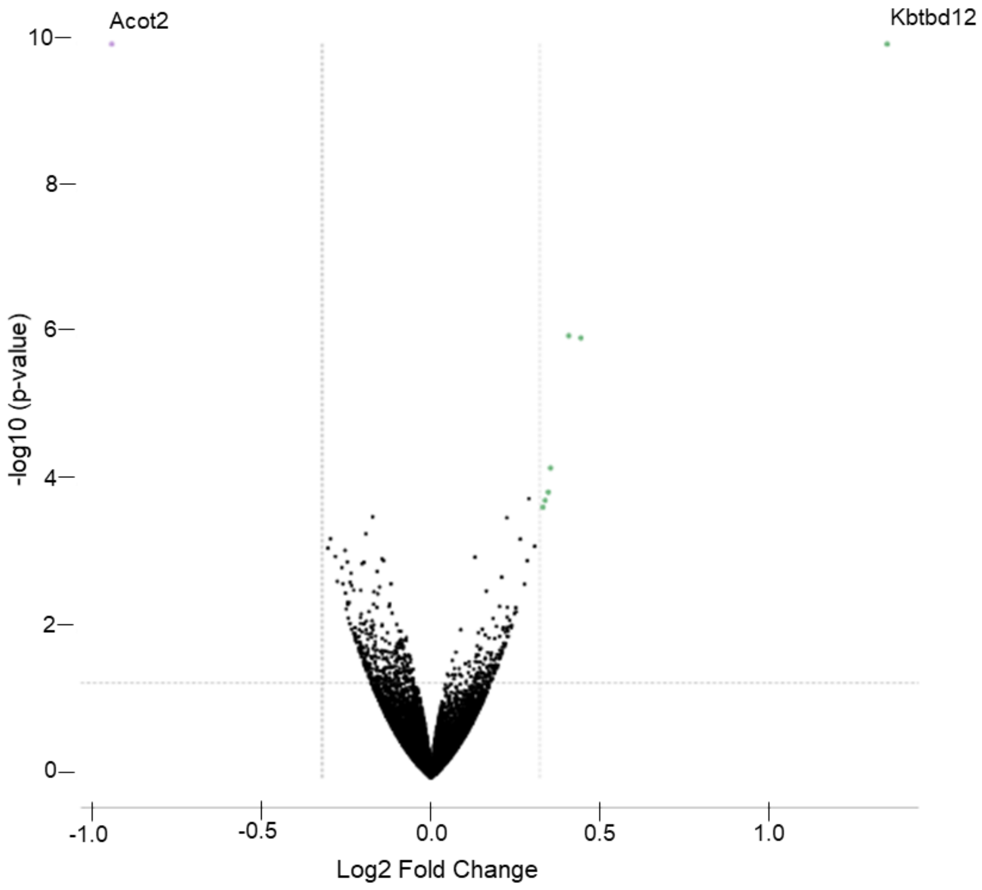


WikiPathways
Focal Adhesion-PI3K-Akt-mTOR-signaling pathway: p-adj 0.00651

| **Genes** | **Fold change** |
|---|---|
| THBS4 | 1.326 |
| FLT4 | 1.278 |
| RELN | 1.271 |

### Supplemental Figure 3

#### B. Fasted FL/FL vs. Acot2-/-

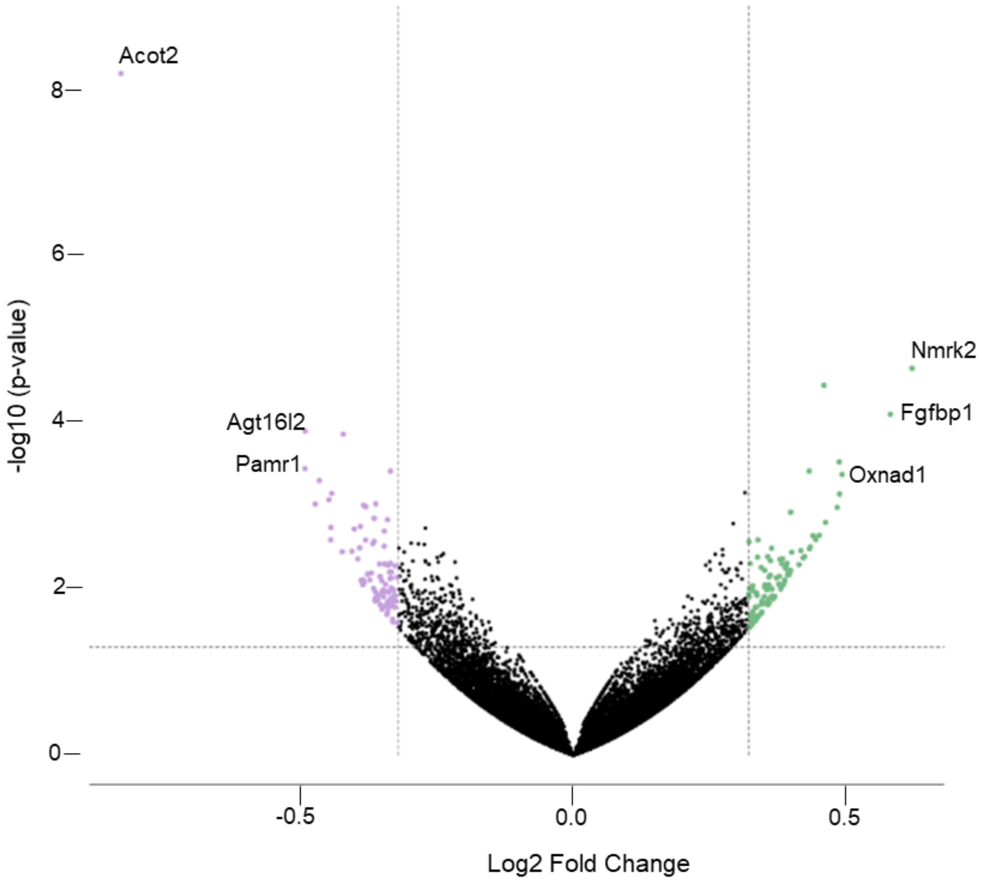


WikiPathways
Microglia Pathogen Phagocytosis Pathway: p-adj 0.02725

| Genes | Fold change |
|---|---|
| C1QA | 1.359 |
| FCER1G | 1.301 |
| PIK3CG | 1.282 |
| NCKAP1L | 1.281 |
| C1QC | 1.275 |

Supplemental Figure 3

C. HFD FL/FL vs. Acot2-/-

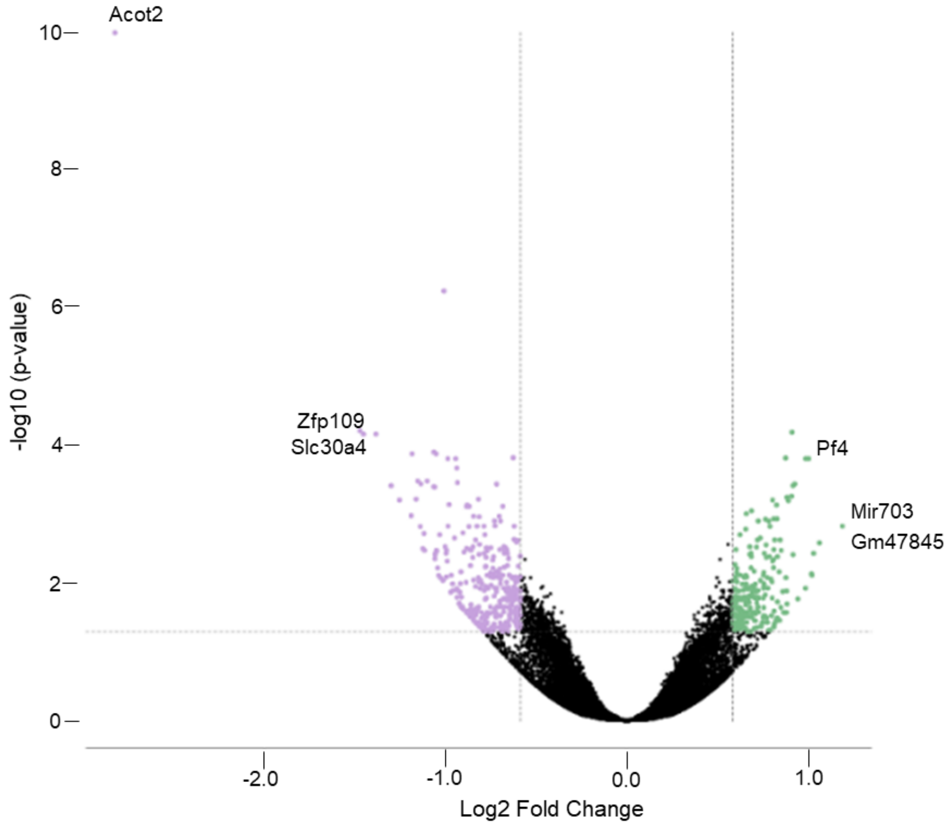


WikiPathways

Electron Transport Chain: p-adj 4.1e-09

| Genes | Fold change |
|---|---|
| *Complex I* | |
| ND5 | -1.835 |
| NDUFB9 | 1.734 |
| NDUFB7 | 1.664 |
| NDUFC2 | 1.657 |
| NDUFB5 | 1.633 |
| NDUFA5 | 1.587 |
| NDUFS3 | 1.539 |

| Genes | Fold Change |
|---|---|
| *Complex III* | |
| UQCRH | 1.641 |
| UQCRB | 1.571 |

| Genes | Fold Change |
|---|---|
| *Complex IV* | |
| COX1 | -1.992 |
| COX6A1 | 1.673 |
| COX6B1 | 1.628 |
| COX4I1 | 1.554 |
| COX7A1 | 1.548 |

| Genes | Fold Change |
|---|---|
| *Complex V* | |
| ATP8 | -1.849 |
| ATP5L | 1.742 |
| ATP5H | 1.714 |
| ATP5G3 | 1.681 |
| ATP5O | 1.545 |
| ATP5K | 1.512 |

REACTOME

Synthesis of PIPs at the Golgi membrane: p-adj 0.01943

| Genes | Fold change |
|---|---|
| PIK3C2A | -1.968 |
| INPP5E | -1.918 |
| SACM1L | -1.893 |
| FIG4 | -1.673 |
| PI4KA | -1.643 |

**Suppl. Fig. 3. Transcriptomics analyses: Quad from Acot2-/- and control mice.**

A-C. Transcriptomic analysis. Volcano plots for chow-fed (A), overnight fasted (B) and 7-day HFD-fed (C) conditions in n=4 mice/ genotype. The log2 Fold Change threshold used was 1.25 for chow and fasted conditions and 1.5 for HFD condition. The adjusted p-value threshold was 0.05 for all conditions. Pathway analysis through https://www.rosalind.bio/ using WikiPathways (in chow, fasted, HFD conditions) and Reactome (in HFD condition), shows the pathways with the best p-adj.

Supplemental Figure 4

All panels: Isolated mitochondria from all limb muscles

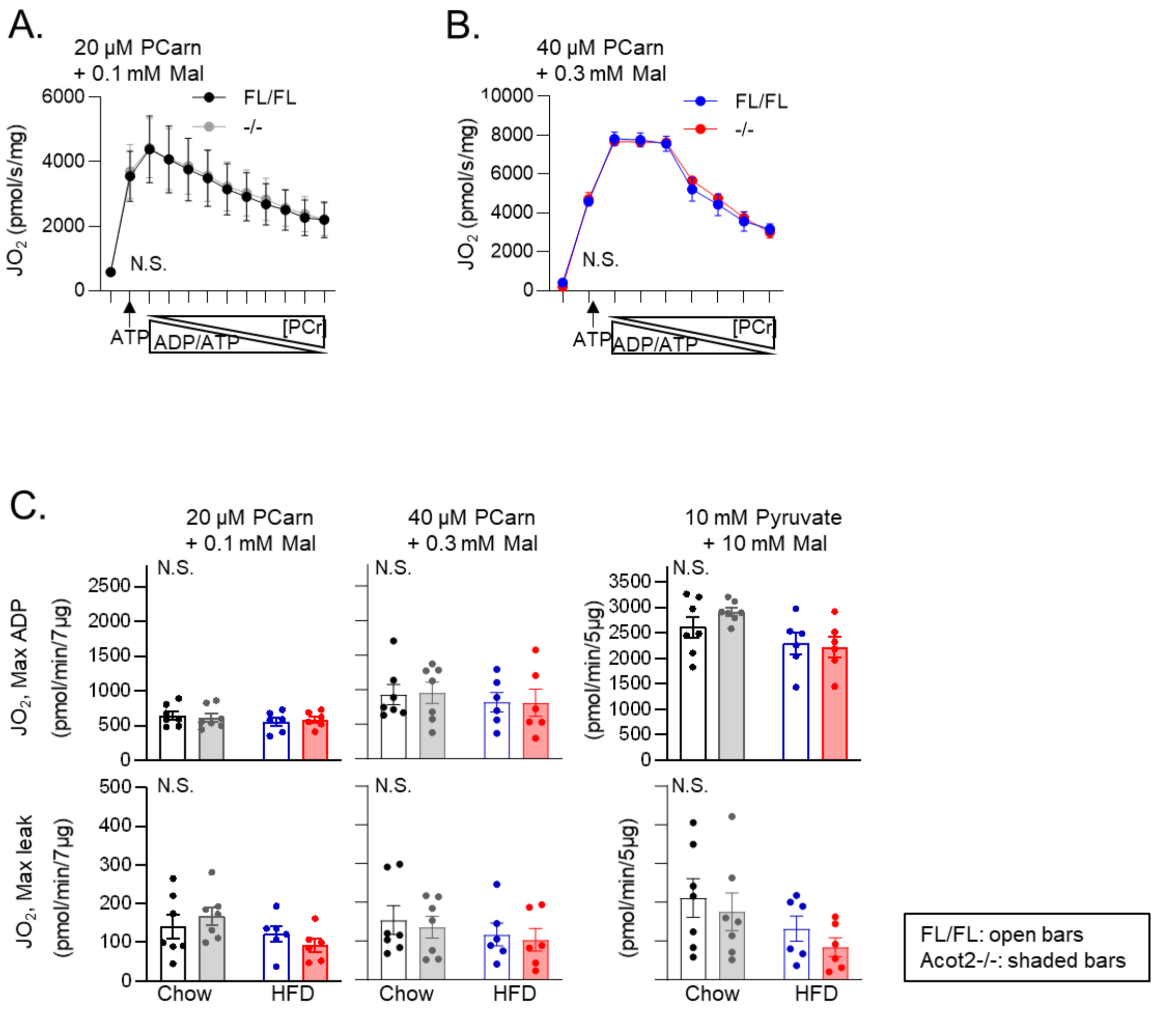

**Suppl. Fig. 4: Bioenergetics analysis in mitochondria isolated from all limb muscles: lack of change with Acot2 loss.**

A. Oxygen consumption ($JO_2$) in mitochondria isolated from all limb muscles of Chow-fed mice, and supplied with 20 µM PCarn and 0.1 mM malate. The creatine kinase clamp approach was used to simulate changes in ATP demand (ADP/ATP) by varying phosphocreatine (PCr) concentration. Values: mean ± s.e.m.. N.S., not significant (unpaired t-test at each step).

B. Same as Panel A, except that mitochondria were isolated from 7-day HFD-fed mice, and 40 uM PCarn + 0.3 mM malate was used. Values: mean ± s.e.m.. N.S., not significant (unpaired t-test at each step).

C. Mitochondrial oxygen consumption ($JO_2$) measured in mitochondria isolated from all limbs muscles of chow- and 7-day HFD-fed mice. The points on the bar graphs represent individual experiments. *Upper*: Measurement of maximal oxidative phosphorylation (max oxphos) in n=7/genotype chow fed mice and n=6/genotype HFD fed mice using different substrates: palmitoyl carnitine (PCarn, 20 µM and 40 µM), and pyruvate (10 mM) in combination with malate (Mal, 0.1 mM, 0.3 mM, 1 mM and 10 mM). *Lower*: Measurement of maximal non-phosphorylating oxidation (max leak) in n=7/genotype chow fed mice and n=6/genotype HFD fed mice using the same substrates as for max oxphos. Bars: mean ± s.e.m. Statistical comparison was by two-way ANOVA and multiple comparisons. N.S. not significant.

### Supplemental Figure 5

A. β-oxidation intermediates in Quad - Pearson's correlation coefficients: lower limit of 65% confidence interval

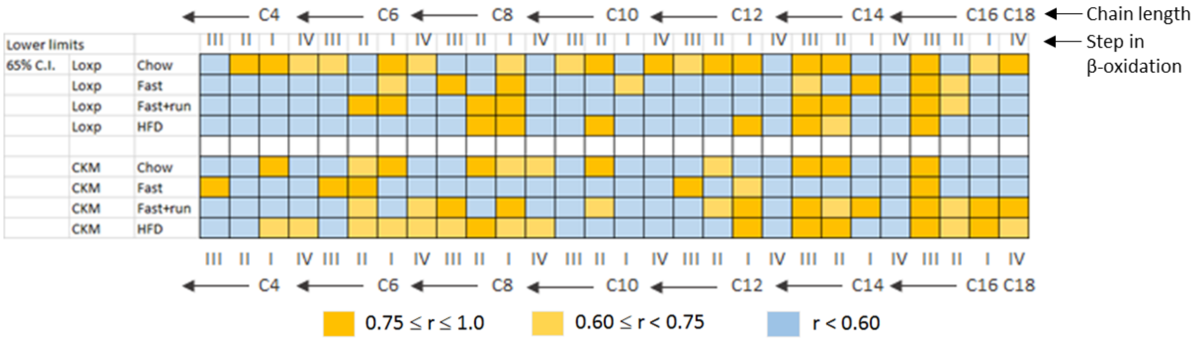


B. **Pearson's correlation coefficients:** q values: pairwise comparisons, BKY, FDR = 5%

| 65% C.I. lower limit | Loxp | CKM |
|---|---|---|
| Chow vs. Fast | 0.01 | 0.35 |
| Chow vs. Fast+run | 0.02 | 0.12 |
| Chow vs. HFD | 0.02 | 0.09 |
| Fast vs. Fast+run | 0.65 | 0.02 |
| Fast vs. HFD | 0.65 | 0.01 |
| Fast+run vs. HFD | 0.74 | 0.3 |

| 65% C.I. lower limit | Loxp vs. CKM |
|---|---|
| Loxp Chow vs. Acot2 Chow | 0.0025 |
| Loxp Chow vs. Acot2 Fast | < 0.001 |
| Loxp Chow vs. Acot Fast+run | 0.12 |
| Loxp Chow vs. Acot2 HFD | 0.2 |
| Loxp Fast vs. Acot2 Fast | 0.15 |
| Loxp Fast vs. Acot2 Fast+run | 0.3 |
| Loxp Fast vs. Acot2 HFD | 0.22 |
| Loxp Fast+run vs. Acot2 Fast+run | 0.36 |
| Loxp Fast+run vs. Acot2 HFD | 0.75 |
| Loxp HFD vs. Acot2 HFD | 0.3 |

**Suppl. Fig. 5. β-oxidation intermediates provide evidence for overload in white SM from Chow-fed and fasted Acot2-/- mice: 65% confidence intervals for Pearson's correlation coefficient analysis**

A. 65% confidence interval (lower limit) for the Pearson's correlation coefficient (r) analysis between adjacent steps of β-oxidation, using the intermediates shown in Fig4B. Roman numerals: enzymes for each step (see Fig4A).

B. Analysis of the 65% confidence interval (lower limit) for the Pearson's correlation coefficients. Statistics: shown within the panel; BKY: Benjamini, Krieger, Yekutieli, to correct for multiple comparisons, FDR: p=0.05.

### Supplemental Figure 6

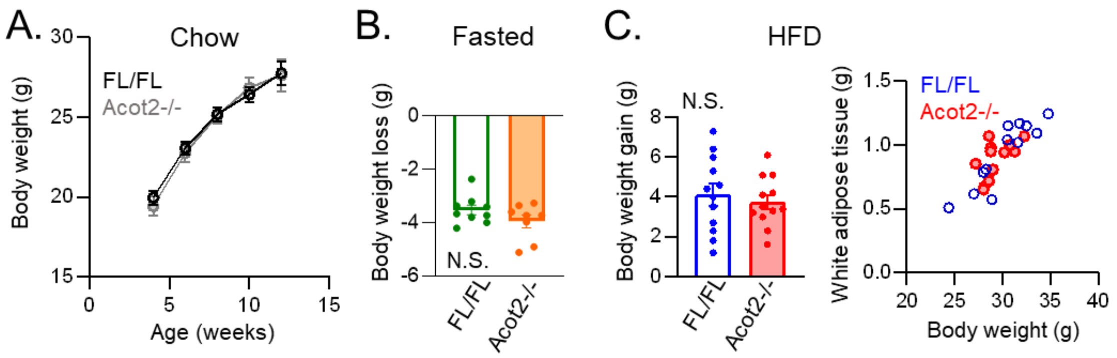

**Suppl. Fig. 6. Lack of change in body weight in mice with Acot2 loss in striated muscle, regardless of feeding regime**

A. Body weight monitored in chow fed mice age 4 to 14 weeks in n=16 FL/FL, n=15 Acot2-/- mice.

B. Weight loss measured in overnight (16 hrs) fasted mice, n=9 FL/FL, n=8 Acot2-/-.

C. Left: Weight gain measured in 7-day HFD (high fat diet) fed mice in n=12/genotype. Right: Weight of white adipose tissue *versus* body weight in 7-day HFD-fed mice, n=12/genotype.

Supplemental Figure 7

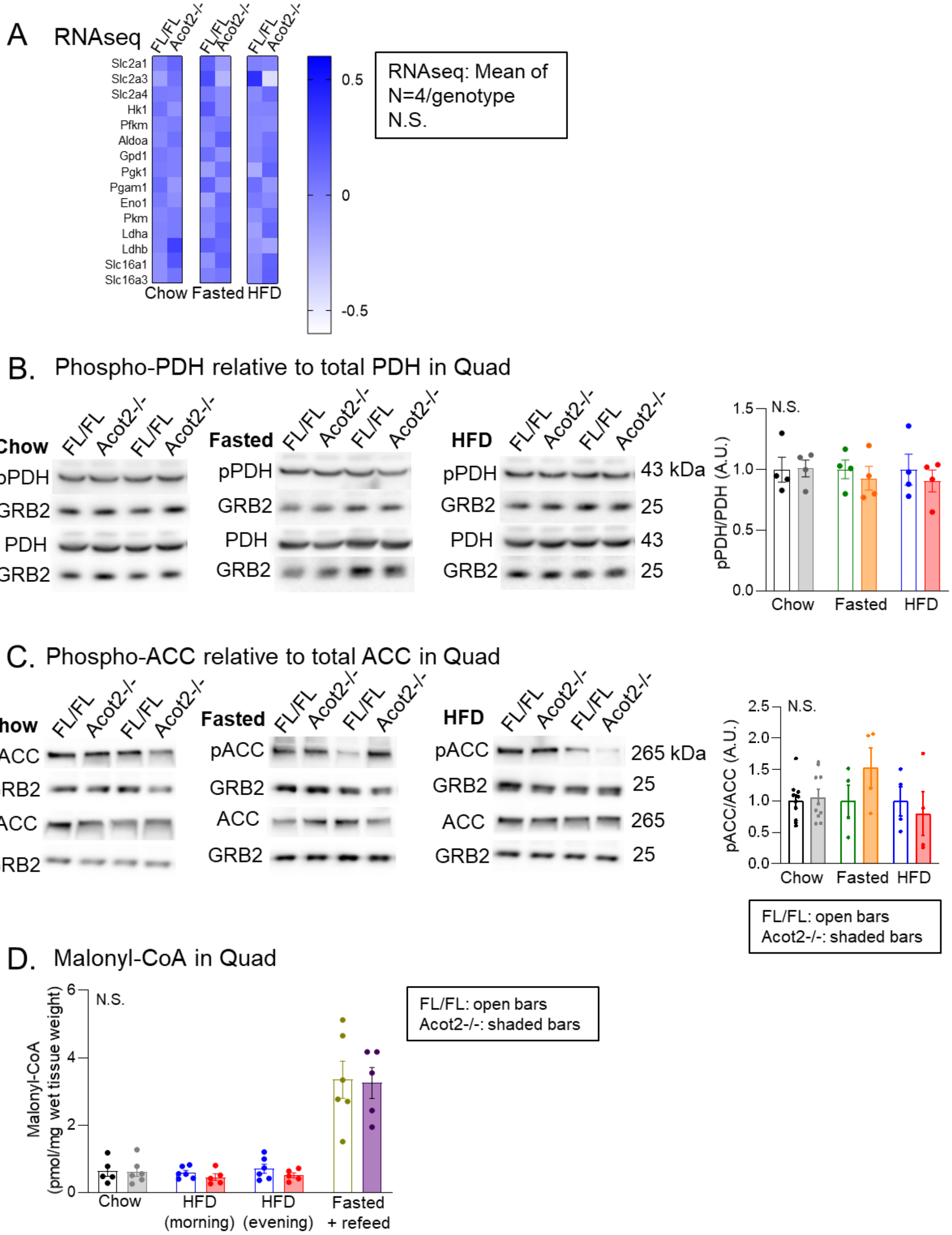

**Suppl. Fig. 7. Lack of effect of Acot2 loss in striated muscle on transcripts for glucose transporters and enzymes, and on pPDH, pACC and malonyl-CoA levels in Quad**

A. RNAseq analysis of Quad muscle, from mice fed normal chow, overnight fasted, or fed HFD for 7 days. Shown are transcript levels for enzymes and transporters involved in glucose transport and metabolism.

B. Representative immunoblots comparing pPDH relative to PDH levels in FL/FL and Acot2-/- quadriceps lysates and GRB2 as loading control. Chow, fasted and HFD conditions. Quantification in n=4/genotype.

C. Representative immunoblots comparing pACC relative to ACC levels in FL/FL and Acot2-/- quadriceps lysates and GRB2 as loading control. Chow, fasted and HFD conditions. Quantification in n=4/genotype.

D. Comparison of malonyl-CoA abundance by mass spectrometry of quadriceps samples collected from FL/FL and Acot2-/- mice studied under chow, HFD morning harvest, HFD evening harvest and refeed (for 6 h) after fasting (for 24 h) conditions in n=6/genotype.

Panel B-C: values are mean ± s.e.m. Statistical comparison was by unpaired t-test. N.S.: not significant. The points on the bar graphs represent individual experiments. N.S. Not significant, two-way ANOVA.

### Supplemental Figure 8

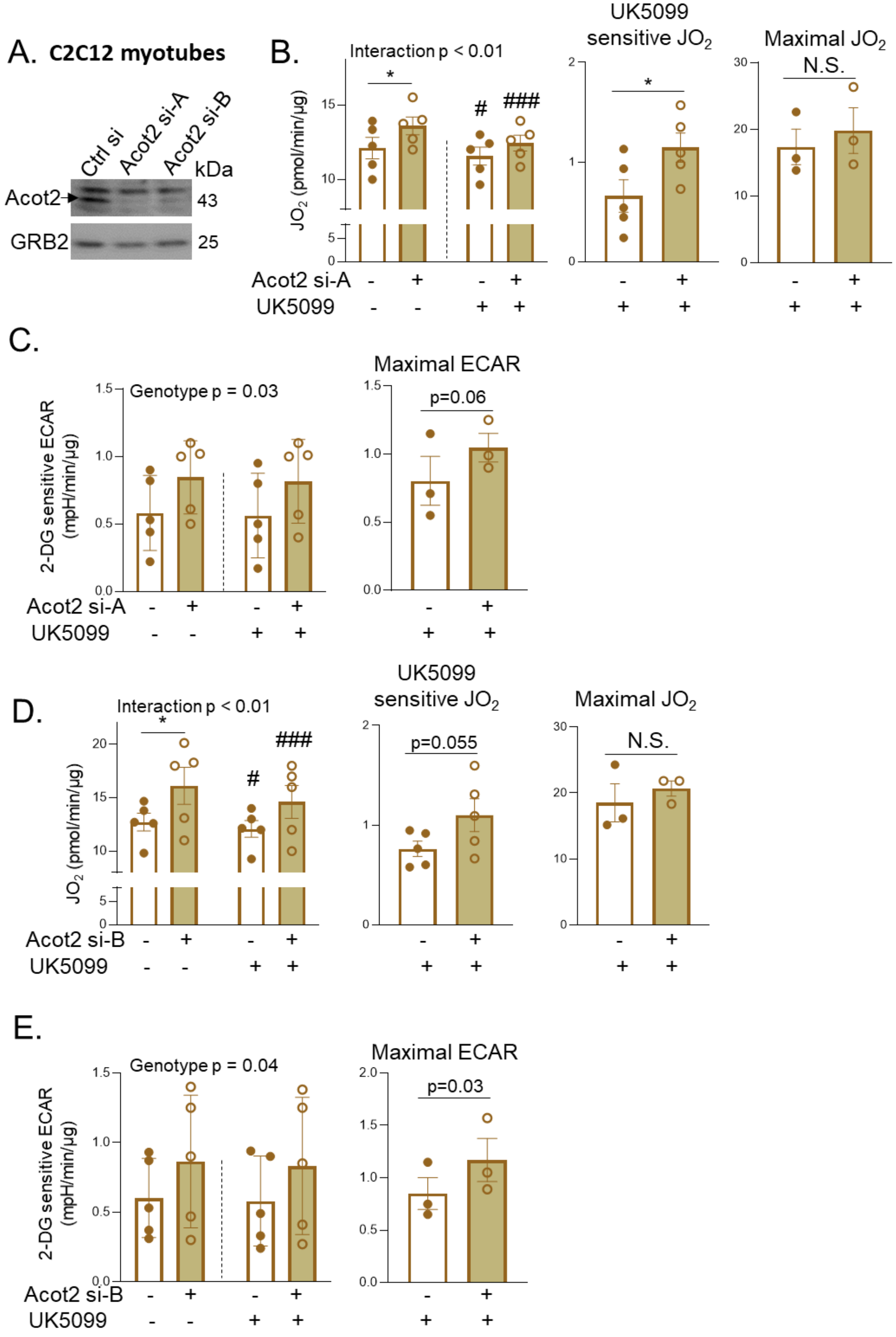

### Supplemental Figure 8

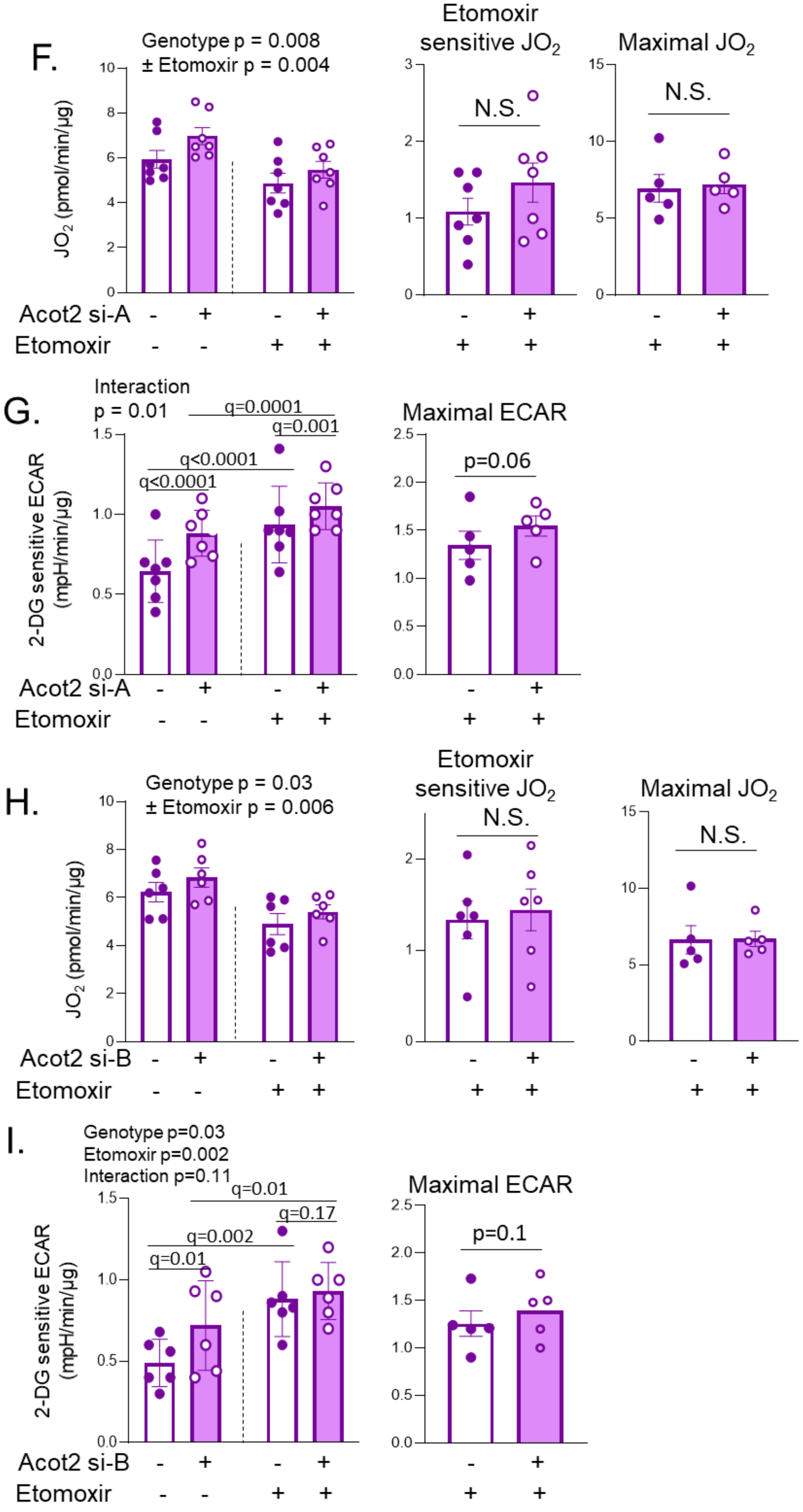

**Suppl. Fig. 8. Increased glucose oxidation in C2C12 myotubes depleted of Acot2**

A. Representative immunoblots comparing Acot2 levels in C2C12 myotubes with addition of 10 pmol control siRNA, Acot2 siRNA-A and siRNA-B and GRB2 as loading control.

B, D. Oxygen consumption ($JO_2$) measured in C2C12 control and Acot2 KD (with addition of 10 pmol siRNA-A or siRNA-B) myotubes, in n=5/condition. Substrates: 10 mM glucose, then 100 µM palmitate:BSA (6:1) in FAO assay medium was added just before the run. Left: basal oxphos before and after 2 µM UK5099 injection to inhibit the mitochondrial pyruvate carrier. Middle: Calculated UK5099 sensitive $JO_2$ in n=5/condition. Right: Maximal uncoupled (FCCP-driven injected after oligomycin injection) $JO_2$, in n=3/condition.

C, E. Extracellular acidification rate (ECAR) measured in C2C12 control and Acot2 KD (with addition of 10 pmol siRNA-A or siRNA-B) myotubes, in n=5/condition, and in the same substrate conditions as in B, D. Left: 2-DG sensitive ECAR before and after 2 µM UK5099 injection. Right: ECAR measured after oligomycin+FCCP injection, in n=3/condition.

F, H. Oxygen consumption ($JO_2$) measured in C2C12 control and Acot2 KD (with addition of 10 pmol siRNA-A or siRNA-B) myotubes, in n=7/condition (siRNA-A) or n=6/condition (siRNA-B). Substrates: 10 mM glucose, then 100 µM palmitate:BSA (6:1) in FAO assay medium was added just before the run. Left: basal oxphos before and after 4 µM Etomoxir injection to inhibit CPT1. Middle: Calculated Etomoxir sensitive $JO_2$. Right: Maximal uncoupled (FCCP-driven injected after oligomycin injection) $JO_2$, in n=5/condition.

G, I. Extracellular acidification rate (ECAR) measured in C2C12 control and Acot2 KD (with addition of 10 pmol siRNA-A or siRNA-B) myotubes, in n=7/condition (si-RNA-A) or n=6/condition (siRNA-B), under the same conditions as in F, H. Left: 2-DG sensitive ECAR before and after 4 µM Etomoxir injection. Right: ECAR measured after oligomycin+FCCP injection, in n=5/condition.

Panels B-I: Bars are the mean ± s.e.m, and points are data from separately cultured flasks of cells tested on separate Seahorse plates. Statistical comparison was by two-way ANOVA (p values) and multiple comparisons (symbols values) with correction for FDR (0.05; Benjamini, Krieger, Yekutieli method) or, when comparison was only between genotypes, by paired t-test (p value). N.S.: not significant.

### Supplemental Figure 9

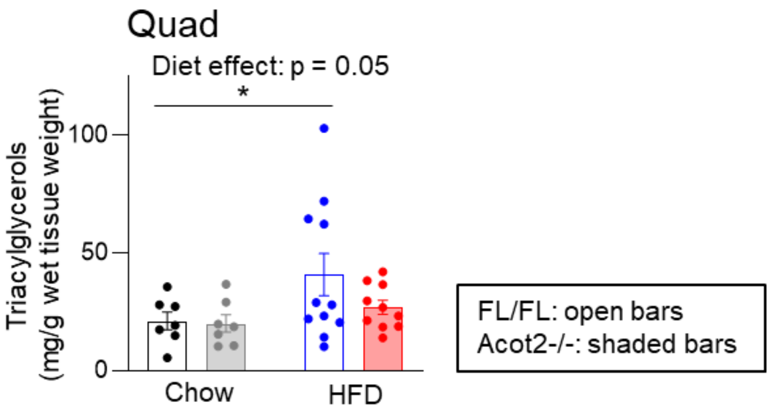


**Suppl. Fig. 9. Lack of significant effect of Acot2 depletion on triacylglycerides in Quad**

Triacylglycerides measured biochemically in Quad from Chow and 7-day HFD-fed mice, n=7-11/genotype. Values: Bars are mean ± s.e.m, points are individual data from each mouse. Statistics: two-way ANOVA (p value).
